# Diet and breeding productivity in European Shag (*Gulosus aristotelis*): insights from two Portuguese colonies

**DOI:** 10.64898/2026.03.29.715095

**Authors:** Bruna Vieira, David Gonçalves, Nuno Oliveira

## Abstract

Climate change and anthropogenic pressures are reshaping marine food webs, altering prey availability and affecting top predators. The European Shag (*Gulosus aristotelis*), a coastal demersal seabird, provides a valuable model for examining environmentally mediated dietary variation, given its trophic plasticity and capacity to adjust prey use according to local availability, while also allowing assessment of potential demographic consequences. This study investigated spatial and temporal variation in diet at two Portuguese colonies (Berlengas and Arrábida) between 2016 and 2024 and assessed long-term reproductive productivity at Berlengas. A total of 468 regurgitated pellets were analysed, and diet composition was quantified using the Index of Relative Importance (IRI). Generalised additive models were applied to assess environmental, spatial, and period-specific effects on diet composition, while reproductive productivity was modelled in relation to prey biomass. Diet variation was primarily explained by environmental predictors, including sea surface temperature, chlorophyll-a concentration, and zooplankton, whereas year per se had no significant effect, indicating environmentally mediated bottom-up effects. Spatial differences between colonies reflected contrasting prey field structures, and period-specific patterns suggested increased specialisation during breeding. Higher biomass of sandeels (Ammodytidae) was positively associated with reproductive output, whereas shifts toward lower-energy prey were associated with reduced productivity. These findings demonstrate that environmentally driven dietary change has measurable demographic consequences, underscoring the importance of bottom-up processes in shaping seabird population dynamics and informing conservation strategies under ongoing climate change.

## Introduction

Cumulative pressures such as global warming, habitat degradation, overfishing, pollution, and biological invasions are reshaping marine ecosystems by altering key abiotic conditions and reducing species resilience (Gladstone-Gallagher et al., 2023; Simeoni et al., 2023). These changes are associated with shifts in species distributions, community structure, and trophic interactions, in some cases leading to substantial reorganisation of marine food webs (Ullah et al., 2018; Ammar et al., 2025). Seabirds occupy upper trophic levels and function as sensitive indicators of marine ecosystem dynamics. Their population trends, breeding success, and foraging behaviour often reflect changes in prey availability and structure (Furness & Tasker, 1999; Piatt et al., 2007). Because primary productivity underpins marine food webs, fluctuations in oceanographic conditions can propagate through higher trophic levels, influencing forage fish and ultimately affecting seabird diet and demography (Gagnon et al., 2013; Perkins et al., 2018). Consequently, seabird diet and population dynamics are strongly influenced by bottom-up processes (Gagnon et al., 2013; Perkins et al., 2018), underscoring the importance of seabirds in sustaining the structural integrity and functional stability of marine and coastal ecosystems (Rajpar et al., 2018). Variability in environmental drivers such as temperature can therefore modify prey availability and spatial distribution through bottom-up processes, leading to changes in predator foraging strategies (Franks, 1992; Heath et al., 2014). Climate-driven trophic effects have been widely documented. For example, breeding failures in Peruvian seabirds followed the collapse of anchoveta stocks, while declines in sandeel populations in the North Sea were associated with reduced breeding success in black-legged kittiwakes (Piatt et al., 2007). Such examples illustrate how environmental variability and prey dynamics can directly influence seabird reproductive performance. Cormorants (Phalacrocoracidae) are highly efficient diving predators adapted to exploit demersal and pelagic fish. Their specialised morphology and physiology enable effective underwater pursuit and capture of prey, allowing them to exploit a wide range of resources and occupy key positions within coastal marine food webs (Grémillet et al., 2005). Diet composition can strongly influence reproductive success. For example, Van Guilder & Seefelt (2013) showed that Double-Crested Cormorants in the Beaver Archipelago experienced reduced productivity following a shift toward lower-energy prey, underscoring the tight link between diet quality and population dynamics. More broadly, breeding success in seabirds varies across species and time, often driven by environmental pressures such as human disturbance, predation, pollution, disease, and shifts in prey availability and composition (Jones et al., 2008; Sherley et al., 2012; Barati et al., 2010). Within this group, the European Shag (*Gulosus aristotelis*) is a coastal, demersal forager characterised by strong site fidelity and relatively limited dispersal capacity (Kennedy & Spencer, 2014; Aebischer, 1995). These traits may heighten its vulnerability to localised environmental change, as restricted movement and reduced gene flow can constrain adaptive responses to shifting ecological conditions (Furlan et al., 2012). European Shag diet has historically been dominated by sandeels (Ammodytidae), suggesting periods of marked trophic specialisation (Harris & Wanless, 1993; Hillersøy & Lorentsen, 2012; Howells et al., 2018). However, climate change and fishing pressure have altered prey availability in many regions, and more recent studies report increased dietary breadth and evidence of trophic plasticity (Velando & Freire, 1999; Hillersøy & Lorentsen, 2012). In Portugal, previous research has documented spatial and seasonal variation in diet among colonies, including the Berlengas and Arrábida, where sandeels often dominate during the breeding period but other demersal and pelagic species contribute variably across time and space (Nascimento et al., 2021; Corona, 2021). The European Shag is currently classified as Vulnerable in Portugal (Almeida et al., 2022), and projections indicate rapid environmental change along the Portuguese coast (Espinosa & Portela, 2025). Understanding how diet responds to environmental variability and how this variation influences reproductive success, is therefore critical for assessing population resilience under ongoing climate change. Accordingly, this study examined spatial and temporal variation in European Shag diet in Portugal, focusing on differences between the Berlengas and Arrábida colonies across breeding and non-breeding periods. We further evaluated long-term reproductive patterns at the Berlengas colony to investigate whether environmentally mediated dietary shifts were associated with changes in reproductive productivity.

## Materials and methods

### Study sites

This study was conducted at two of the main European Shag breeding sites in Portugal, representing spatially distinct ecosystems: the insular Berlengas Archipelago (comprising Berlenga Grande, Estelas, and Farilhões islets) and the coastal Professor Luiz Saldanha Marine Park (Arrábida). The Berlengas (39°24′N, 9°30′W), located 10 km off Peniche, is the primary breeding site in the country and is protected as a Natural Reserve, a Special Area of Conservation (SAC), a Special Protection Area (SPA), and an Important Bird Area (IBA) (Nascimento, 2018; BirdLife International, 2025). Arrábida (38°26′N, 9°00′W), part of Arrábida Natural Park, hosts one of the largest national colonies and features structurally complex shallow-water habitats with rocky substrates, kelp forests, and seagrass meadows, providing key feeding and breeding grounds (Corona, 2021). Both sites are designated Marine Protected Areas (MPAs) and play a critical role in species conservation. Figure 1 shows the location of the study areas, Berlenga Grande and Arrábida.

**Figure 1.**
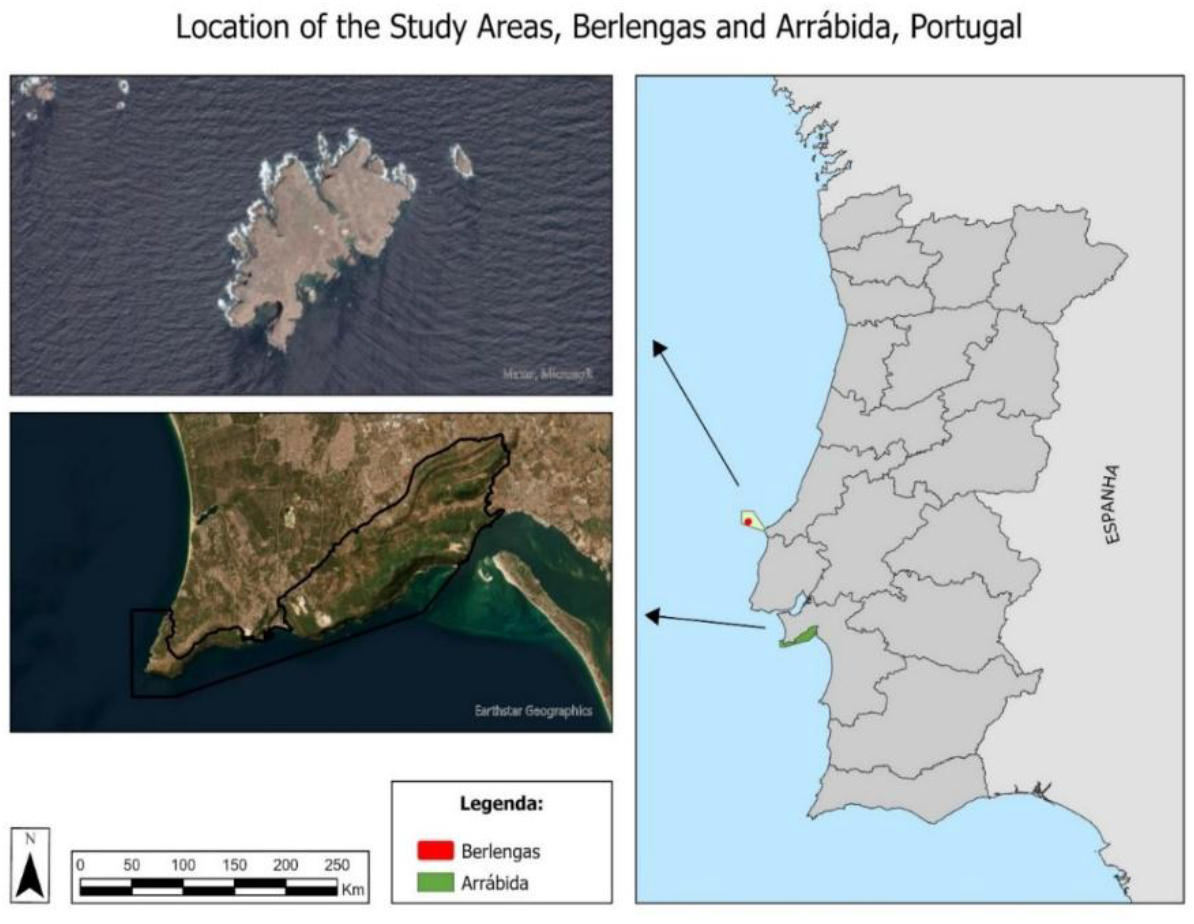
Location of the study areas: Berlenga Grande and Arrábida, Portugal.

### Sample collection

Regurgitated pellets were collected at Berlenga Grande Island (2016-2024) and Arrábida (2020-2021) during both the breeding (February-July) and non-breeding (August-January) periods. Due to limited nest access, pellets were collected from roosting sites, frozen, and individually labelled. A total of 468 pellets were collected, including 375 from the breeding period and 93 from the non-breeding period (Table 1). The unequal sample sizes across periods were taken into account when interpreting seasonal dietary variation.

**Table 1.** Number of regurgitated pellets collected for diet analysis of the European Shag (Gulosus aristotelis) during the breeding and non-breeding periods at the Berlengas (2016-2024) and Arrábida (2020-2021) colonies. Dashes (-) indicate no sapling conducted.

| Year | Berlenga Grande Island |  | Arrábida Coast |  |
| --- | --- | --- | --- | --- |
|  | Breeding | Non-breeding | Breeding | Non-breeding |
| 2016 | 8 | 5 | - | - |
| 2017 | 83 | 23 | - | - |
| 2018 | 9 | 2 | - | - |
| 2019 | 69 | 3 | - | - |
| 2020 | 15 | 7 | - | 44 |
| 2021 | 25 | 2 | 50 | - |
| 2022 | 30 | 6 | - | - |
| 2023 | 59 | 1 | - | - |
| 2024 | 27 | - | - | - |

Productivity monitoring was conducted exclusively at Berlengas (2016-2024) by recording adults, eggs, hatched chicks, and juveniles using telescopes and standardised sheets (see Figure S1 (Online Resource 1). Due to the lack of direct dietary data per nest, productivity was assessed across all monitored nests (n = 400), while diet was characterised from breeding-period pellets (Table 2).

**Table 2.**
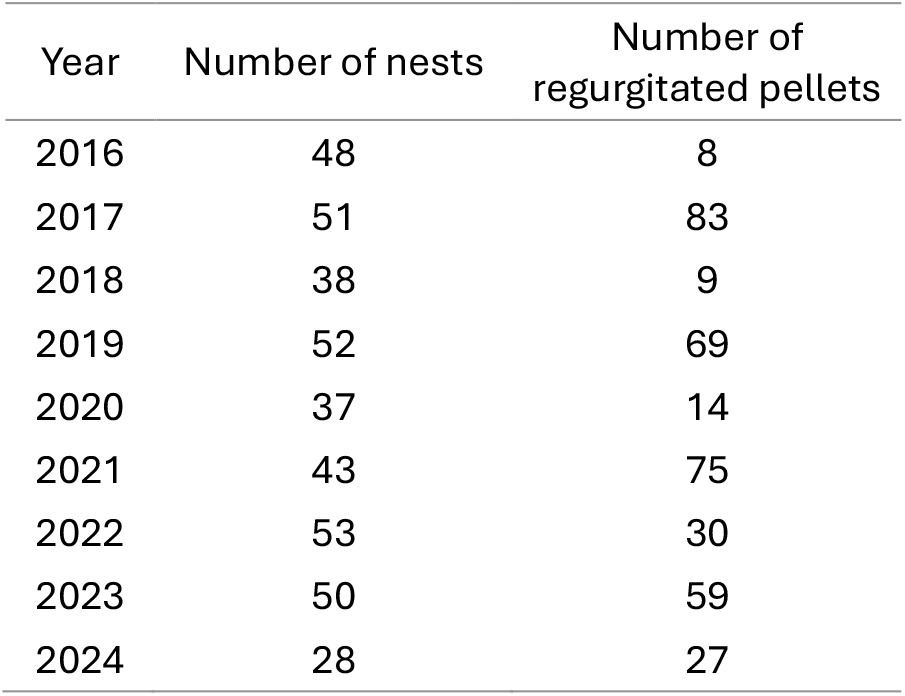
Yearly number of monitored nests (productivity data) and collected regurgitated pellets (diet data) in Berlengas colony, used to evaluate potential links between diet composition and breeding success.

| Year | Number of nests | Number of regurgitated pellets |
| --- | --- | --- |
| 2016 | 48 | 8 |
| 2017 | 51 | 83 |
| 2018 | 38 | 9 |
| 2019 | 52 | 69 |
| 2020 | 37 | 14 |
| 2021 | 43 | 75 |
| 2022 | 53 | 30 |
| 2023 | 50 | 59 |
| 2024 | 28 | 27 |

**Table 3.** The ten most representative prey species/groups selected based on maximum IRI values (2016-2024). Max IRI represents the highest Index of Relative Importance recorded for each species, while Mean IRI ± SD corresponds to the mean IRI and associated standard deviation calculated across the study period (2016-2024).

| Prey species/groups | Max IRI | Mean IRI ± SD |
| --- | --- | --- |
| *Ammodytidae* | 47411.72 | 4169.05 ± 10802.72 |
| *Gaidropsarus mediterraneus* | 6794.45 | 1726.33 ± 3379.05 |
| *Trisopterus luscus* | 10616.20 | 1562.34 ± 3559.65 |
| *Coris julis* | 3653.88 | 808.24 ± 1016.19 |
| *Serranus cabrilla* | 2634.43 | 606.13 ± 1009.22 |
| *Trachurus trachurus* | 1662.72 | 563.05 ± 952.42 |
| *Labrus mixtus* | 3529.41 | 340.76 ± 971.67 |
| *Ciliata mustela* | 1812.86 | 248.09 ± 632.66 |
| *Symphodus bailloni* | 1385.18 | 215.45 ± 433.02 |
| *Atherina presbyter* | 440.16 | 82.67 ± 158.47 |

### Laboratory processing and diet analysis

Pellets were processed following Provencher et al. (2019). Prey identification was based primarily on otoliths, with measurements to the nearest 0.1 mm. Species-level identification was approached conservatively for groups with morphologically similar or eroded otoliths (e.g., Ammodytidae). Fish size and biomass were estimated using published regression equations (see Table S1 (Online Resource 1). Prey species were classified ecologically as demersal, benthopelagic, reef-associated, or pelagic (Froese & Pauly, 2024). Dietary composition was quantified using three complementary metrics (Hart et al., 2002): numerical frequency (%FN = [number of individuals of a species / total number of individuals] × 100), frequency of occurrence (%FO = [number of pellets containing a species / total number of pellets] × 100), and biomass frequency (%FB = [biomass of a given species / total biomass consumed] × 100). To account for biases inherent to each metric, the Index of Relative Importance (IRI, %) was calculated as IRI = %FO × (%FN + %FB) (Pinkas et al., 1971). The ten most representative prey species/groups were selected for simplified presentation. Prey diversity was assessed using the Shannon-Wiener index (H’), enabling quantitative comparisons across periods and areas and integration with productivity analyses.

### Environmental variables

Several environmental variables were used to model diet composition. Chlorophyll-a concentration (chl-a), sea surface temperature (SST) and zooplankton data were obtained from the European Union’s Copernicus Marine Service (Aznar et al., 2016; Jean-Michel et al., 2021), with monthly resolution and a spatial grid of 0.083 degrees (about 9 km). An area corresponding to a buffer of 18 km around Berlenga Grande Island and Peniche Peninsula was defined based on the observed breeding range of European Shag on Berlengas (Nascimento et al., 2023). Values for each variable were averaged by month. A similar approach was conducted for the Arrábida Population. The North Atlantic Oscillation (NAO) index was used as a large-scale climatic proxy influencing oceanographic conditions and prey availability. Monthly values were obtained from NOAA (http://www.cpc.ncep.noaa.gov). NAO mean yearly values were estimated for the period from December to March.

### Statistical analysis

Temporal variation in diet composition was analysed using Generalised Additive Models (GAMs; Banks & Fienberg, 2003), while potential dietary effects on reproductive success were evaluated using Generalised Additive Mixed Models (GAMMs; Ferrari et al., 2025). For diet models, IRI values were modelled as a function of year and environmental predictors. Environmental variables included the NAO, chl-a, SST, zooplankton, and diet diversity (Shannon-Wiener index). Smooth functions were applied to continuous predictors to account for potential non-linear relationships, while season, study area, and species were included as categorical fixed effects. Models assumed a negative binomial error distribution with a log link function. Complementary pairwise comparisons of prey species between colonies (Berlengas vs. Arrábida) were conducted using the non-parametric Mann-Whitney U test applied to IRI values (Mann & Whitney, 1947). Reproductive success at Berlengas was modelled with breeding-period pellets considered sampling units. Explanatory variables included the biomass of the ten most relevant prey species/groups and diet diversity (Shannon-Wiener index per breeding period), with random effects accounting for hierarchical structure and repeated measures. A Tweedie distribution (p ≈ 1.99) with a log link was applied. Significant effects were visualised as described above.

## Results

### Diet composition

A total of 13,418 otoliths were recovered from 467 pellets, of which 5,335 individuals were identified, representing 80 fish species of 58 genera and 29 families. Demersal species dominated the diet (70%), followed by reef-associated (17%) and pelagic species (10%). Average estimated lengths were 12.67 cm, 8.27 cm, and 10.23 cm for demersal, reef-associated, and pelagic species, respectively. During the breeding period, Ammodytidae consistently represented the core prey group, with high values across numerical frequency (%FN), frequency of occurrence (%FO), and biomass frequency (%FB). Non-breeding diets showed higher taxonomic turnover, though Ammodytidae remained recurrent. Overall, %FN and %FO highlighted the stability of key prey groups, whereas %FB indicated interannual shifts in dominance among secondary prey (Table S2, Online Resource 1). The ten most important prey species/groups, in Table 3 (see full table in Table S3, Online Resource 1) showed strong heterogeneity in their Index of Relative Importance (IRI), reflecting pronounced interannual variability. Most species were present throughout the study period, though some occurred sporadically (e.g., *Trachurus trachurus, Gaidropsarus mediterraneus, Atherina presbyter*).

### Drivers of diet variation

The GAM explained 52% of the deviance in IRI values. Significant non-linear effects were observed for SST (χ^2^ (2.13) = 28.29, P < 0.001), chl-a (χ^2^ (1.19) = 13.66, P < 0.001), zooplankton (χ^2^ (1.30) = 15.77, P < 0.001), and the Shannon-Wiener index (χ^2^(1.20) = 21.29, P < 0.001). Year (χ^2^(0.01) = 0.01, P = 0.28) and NAO (χ^2^ (0.00) = 0.00, P = 0.58) were not significant. Parametric coefficients showed significant differences between periods, study areas, and several prey species (P < 0.05). Significant environmental effects on diet composition are illustrated in Fig. 2, while parametric contrasts among categorical predictors are shown in Fig.3.

**Figure 2.**
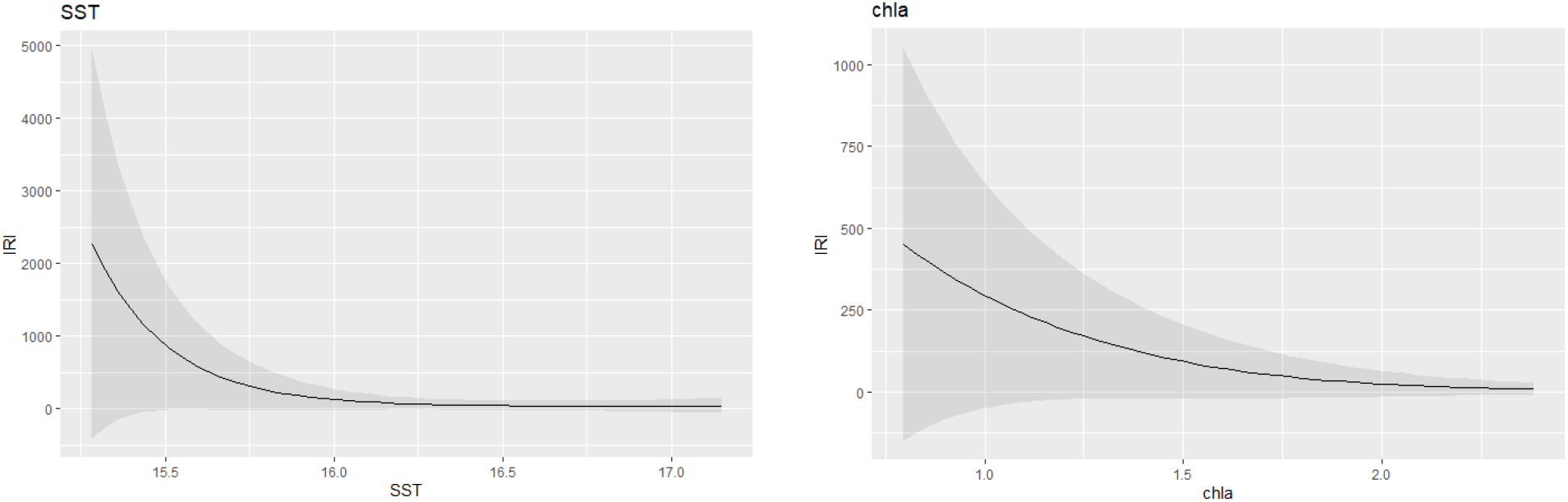
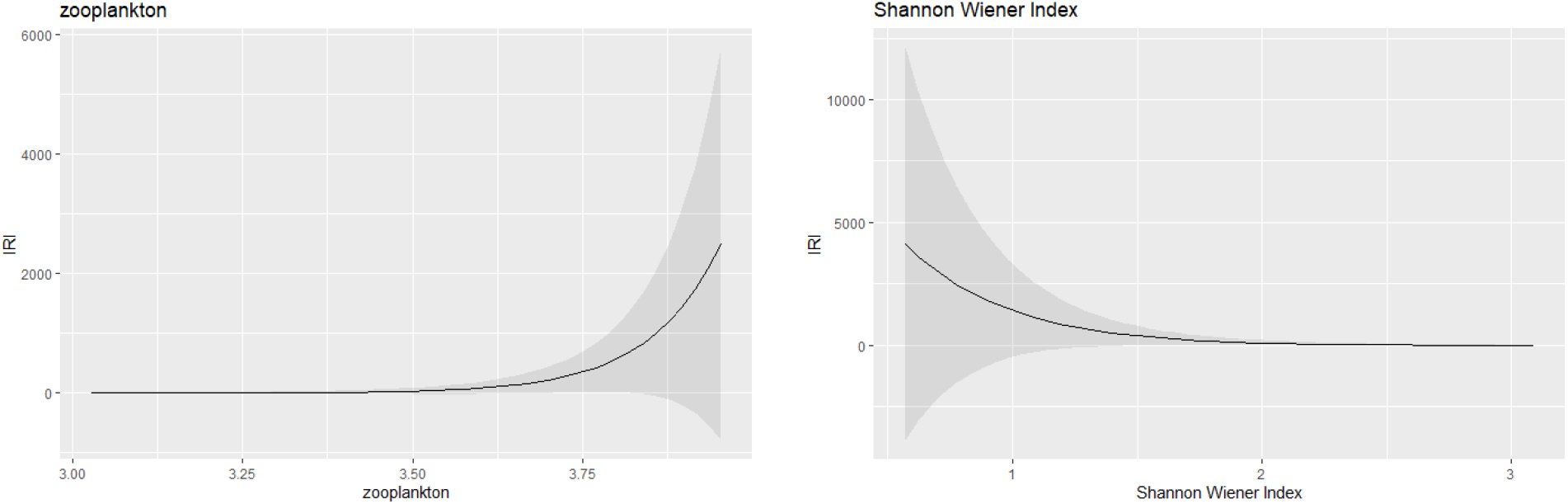
Effects of significant smooth terms from the GAM showing relationships between the Index of Relative Importance (IRI) and sea surface temperature (SST), chlorophyll-a concentration (chla), zooplankton, and Shannon-Wiener diversity index.

IRI was generally higher in Berlengas than Arrábida (Fig. 3), though wide confidence intervals indicated considerable variability among pellets. Ammodytidae consistently had the highest IRI values. Pairwise comparisons in Berlengas revealed significant differences among several prey species (Table S4, Online Resources 1), whereas no such differences were observed in Arrábida.

**Figure 3.**
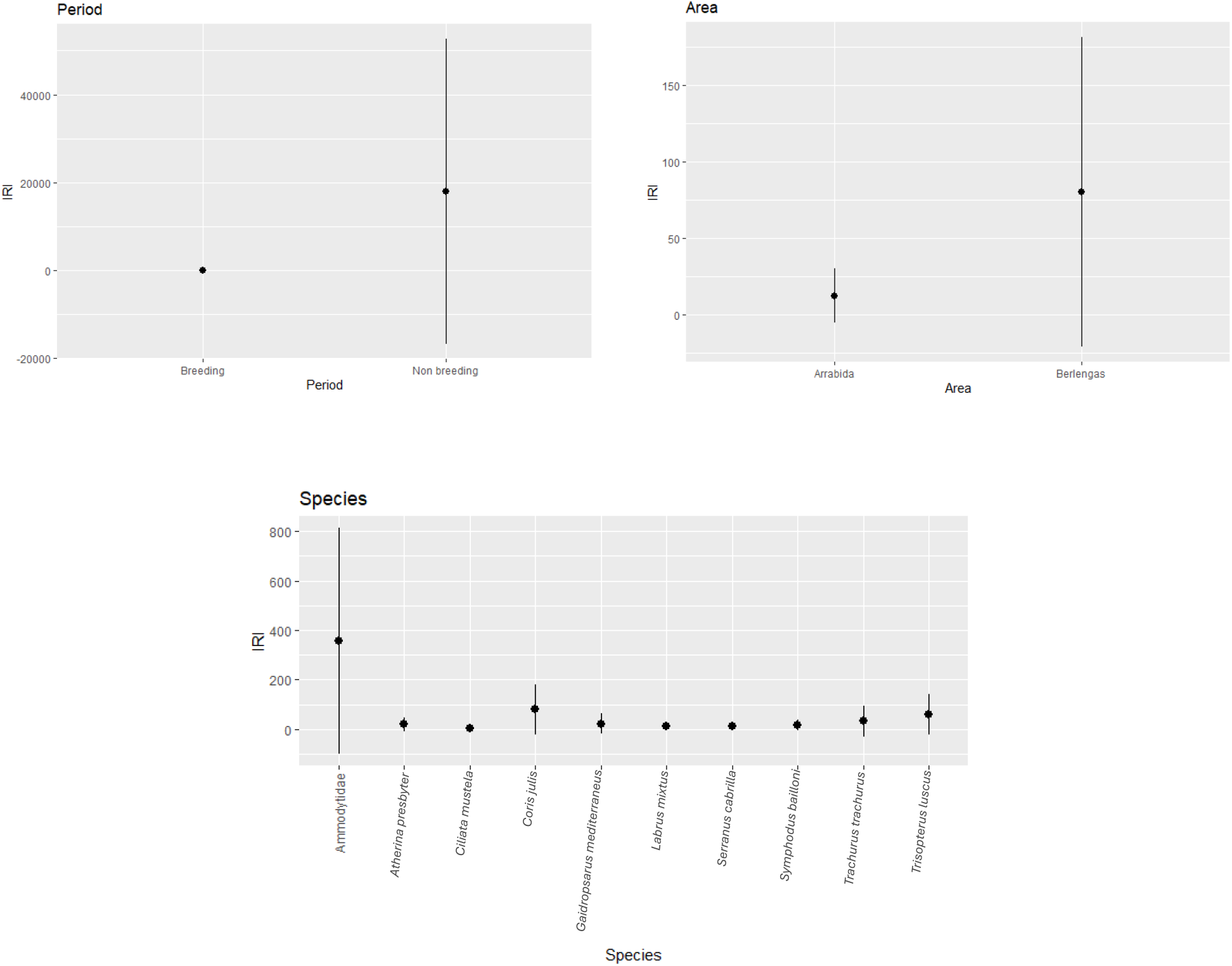
Parametric effects from the GAM showing differences in the Index of Relative Importance (IRI) among period, study areas, and species.

### Productivity analysis (Berlengas only)

Reproductive success was explained by 81% of deviance in the GAMM (adjusted R^2^ = 0.81) and was significantly influenced by Year (F1 = 856.21, P < 0.001) as well as by several of the top 10 species. Significant effects were found for Ammodytidae (F3 = 7.91, P < 0.001), *Labrus mixtus* (F3 = 4.00, P < 0.001), *Trachurus trachurus* (F3 = 3.16, P < 0.001), *Gaidropsarus mediterraneus* (F3 = 1.52, P < 0.05), and *Atherina presbyter* (F3 = 1.81, P < 0.05). The other species were retained but with no statistically significant effect (all p > 0.05). The biomass of *Trachurus trachurus, Labrus mixtus, Gaidropsarus mediterraneus*, and *Atherina presbyter* exhibited a subtle decreasing effect on reproductive success over time (Figures 4), whereas Ammodytidae biomass showed a slight increasing trend (Figure 5). Overall, reproductive success displayed a marked decline throughout the study period (Figure 6).

**Figure 4.**
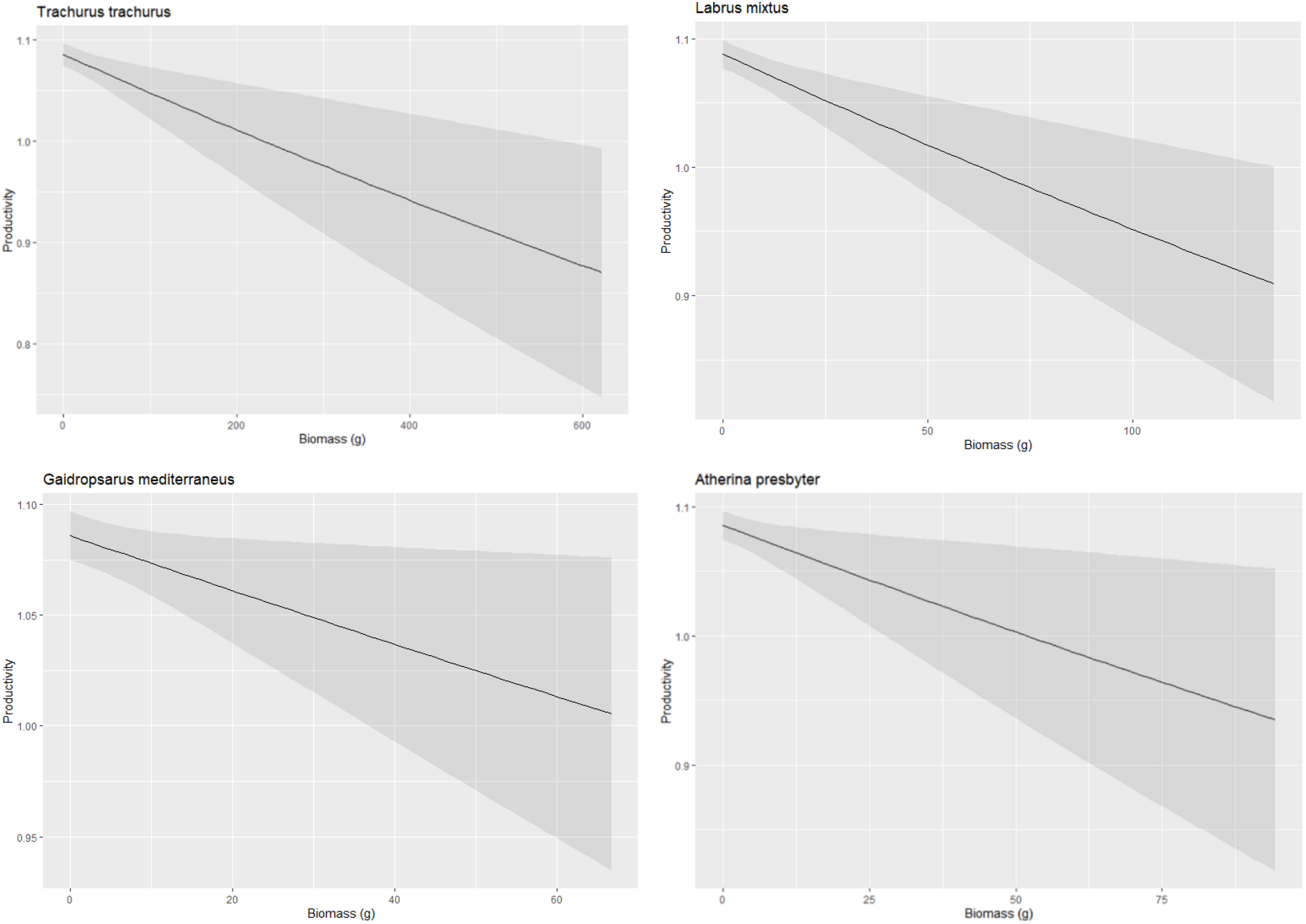
Effect of prey biomass on European Shag reproductive success (productivity): Trachurus trachurus; Labrus mixtus; Gaidropsarus mediterraneus; Atherina presbyter.

**Figure 5.**
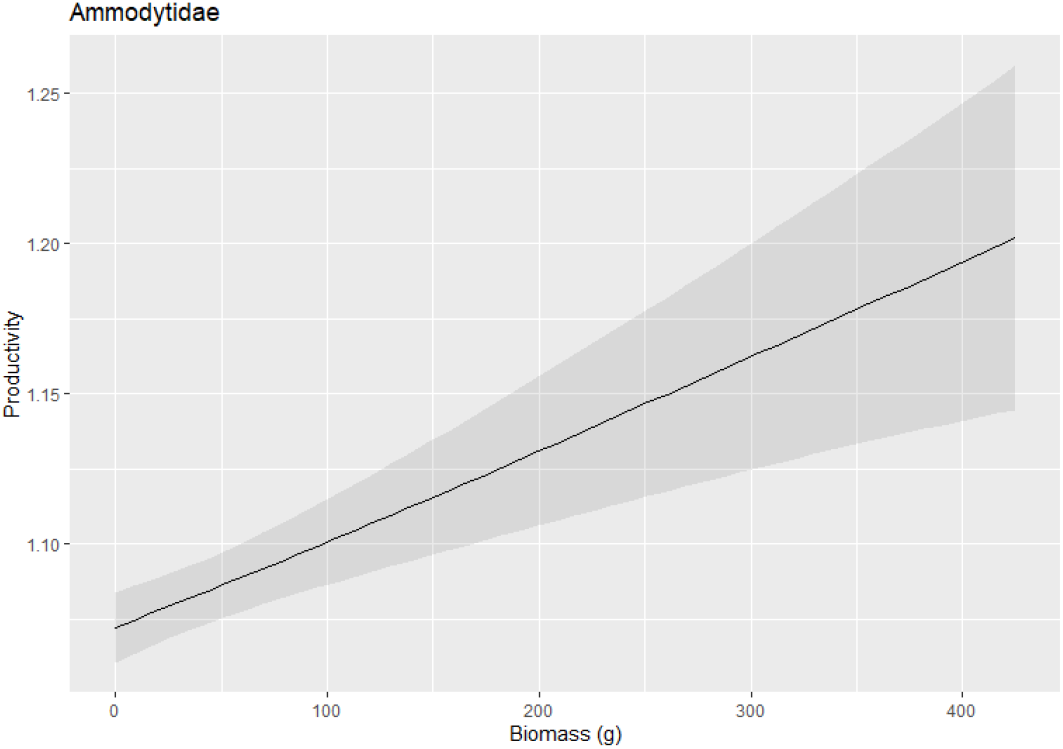
Effect of Ammodytidae biomass on European Shag reproductive success (Productivity).

**Figure 6.**
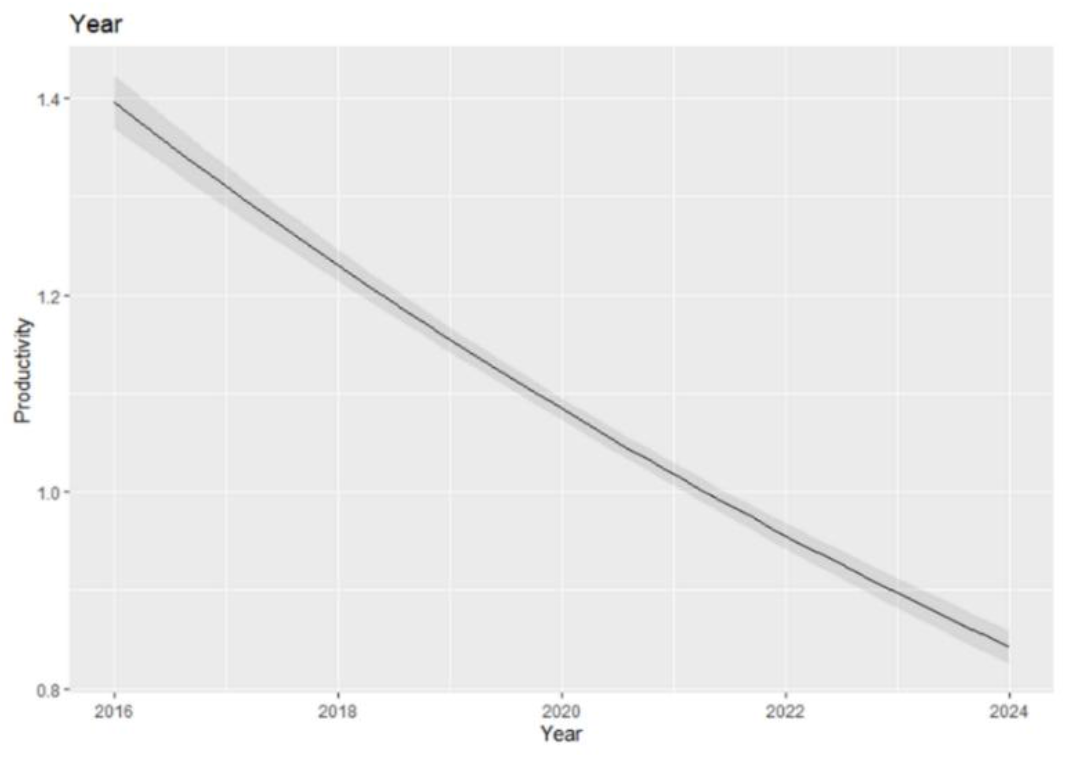
Effect of biomass on Year in Productivity.

## Discussion

Our results demonstrate that diet composition in the European Shag is shaped by environmental variability, spatial context, and life-history constraints. Although the species shows marked trophic plasticity, preferential reliance on energy-rich prey emerged as a key determinant of reproductive performance. Temporal variation in diet was environmentally mediated rather than driven by year per se, while spatial and period-specific differences reflected local prey field structure and central-place foraging constraints. Importantly, shifts away from dominant, energetically profitable prey were associated with reduced productivity, highlighting the demographic consequences of trophic change.

### Dietary diversity of the European Shag

Ammodytidae emerged as the dominant prey group in the European Shag diet, consistent with numerous previous studies (Howells et al., 2018; Harris & Wanless, 1997; Velandol & Freire, 1999). This family represents an energetically optimal resource due to its high abundance and availability (Mitchell et al., 1998), predictable diel behaviours (e.g. vertical larval migrations and adult shoaling during feeding; Winslade, 1974), and elevated lipid content (Staudinger et al., 2020). Such characteristics render sandeels a highly profitable prey, capable of meeting the increased energetic demands during the breeding period. Pout Whiting (*Trisopterus luscus*) constituted the second most important prey species, contributing substantially to total biomass despite relatively low numerical occurrence. This pattern is consistent with a previous study by Velando & Freire (1999), in which the species accounted for only ∼5% of prey items numerically but more than 16% of total biomass. This disproportionate biomass contribution underscores the importance of prey size and energetic value in shaping diet composition. Together, these findings indicate a clear preference for high-energy prey capable of maximising energy intake efficiency. Secondary prey, including Mediterranean Rockling (*Gaidropsarus mediterraneus*), Horse Mackerel (*Trachurus trachurus*), and Mediterranean Rainbow Wrasse (*Coris julis*), contributed moderately to the diet, reflecting consistent but non-dominant exploitation. Horse Mackerel occurred in low numbers, supporting its classification as a minor prey item (Xirouchakis et al., 2017), whereas Mediterranean Rainbow Wrasse, although more frequent, contributed relatively little biomass due to its small size (Al-Ismail et al., 2013; Morat et al., 2014). These species are generally less predictable and energetically variable compared to primary prey, suggesting that their inclusion reflects opportunistic foraging under fluctuating prey availability. Importantly, the relationship between dietary diversity and prey dominance (using IRI in this study as a proxy) showed a significant negative trend, indicating that periods of higher Shannon-Wiener diversity corresponded to reduced dominance by any single species. This pattern supports a dynamic specialist-generalist continuum, whereby European Shags exhibit specialisation when energetically profitable prey are abundant but shift towards a more generalised diet when dominant prey availability declines. Such foraging plasticity likely represents an adaptive response to environmental variability, which allows individuals to buffer fluctuation in prey supply while maintaining energetic intake.

### Environmental drivers of temporal diet variation

Seabirds are widely recognized as sensitive indicators of marine ecosystem dynamics, as fluctuations in their distribution, abundance, and breeding biology often reflect changes in prey availability and structure (Piatt et al., 2007; Furness & Tasker, 1999). Consistent with this, variation in European Shag diet composition was evident across the study period, although year itself did not exert a significant effect on IRI, indicating that temporal changes were not driven by intrinsic interannual trends alone but were instead environmentally mediated. Environmental predictors, including sea surface temperature (SST), chlorophyll-a concentration (chl-a), and zooplankton biomass, explained a substantial proportion of the variation in diet composition, suggesting the influence of bottom-up processes operating through both primary and secondary production. IRI decreased with increasing SST. Elevated sea temperatures can enhance water-column stratification, altering thermal structure and reducing vertical mixing. Such stratification may modify the vertical and horizontal distribution of forage fish, which often avoid abrupt temperature gradients or remain below shallow thermoclines, thereby reducing accessibility to diving seabirds without necessarily decreasing overall productivity (Weeks et al., 2013; Abookire & Piatt, 2005). Interestingly, the North Atlantic Oscillation (NAO) did not significantly affect diet composition. This suggests that local oceanographic conditions, such as SST, may exert more immediate and ecologically relevant influences on prey accessibility than broad-scale atmospheric oscillations. Similar patterns have been documented in other marine systems, where prey availability and seabird demographic responses were more closely linked to regional sea surface temperature variability than to large-scale climatic indices such as El Niño or the NAO, highlighting the stronger ecological relevance of local thermal conditions (Weeks et al., 2013; Sandvik et al., 2004). Chlorophyll-a, widely used as a proxy for primary productivity (Pallavi et al.,2024), was negatively associated with IRI. Increased primary productivity can enhance overall prey availability and community complexity through bottom-up processes (Fernández et al., 2016). Under such conditions, predators may exploit a broader suite of prey taxa, reducing reliance on any single dominant species and leading to lower IRI values. In this sense, elevated primary production may promote dietary diversification rather than specialisation. In contrast, zooplankton biomass (secondary production) showed a positive relationship with IRI. Zooplankton constitutes a critical link between primary production and higher trophic levels, serving as the principal food source for many forage fish species (Vereshchaka, 2024; Ratnarajah et al., 2023). Variability in plankton communities has been linked to interannual fluctuations in fish recruitment in temperate shelf seas, including key forage species such as sandeels, whose early life stages are strongly dependent on zooplankton availability (van der Kooij et al., 2008; Lomartire et al., 2021). Increased secondary production may therefore enhance sandeel recruitment and local aggregation, promoting their dominance in predator diets. Under such conditions, reliance on this energetically profitable prey may increase, resulting in higher IRI values and greater dietary specialisation. Therefore, the contrasting relationships observed for chlorophyll-a and zooplankton biomass suggest that productivity at different trophic levels influences diet composition through distinct mechanisms. While primary productivity represents the potential energy input to the system and may increase overall prey diversity, secondary production more directly determines the recruitment and aggregation of key forage fish species, thereby shaping prey dominance patterns within predator diets.

### Spatial and period-specific diet patterns

Diet composition differed markedly between the two contrasting regions along the Portuguese coast, with higher mean IRI values in Berlengas than in Arrábida, indicating a stronger dominance by key prey species. In Berlengas, Ammodytidae dominated the diet, with intermediate contributions from Pout whiting and Mediterranean rainbow wrasse, while other species were consumed sporadically. This pattern reflects reliance on a few predictable and aggregated prey types, a characteristic commonly associated with insular seabird colonies, as reported for European Shag on Norwegian islands (Barrett et al., 1990). In contrast, diet composition in Arrábida was more evenly distributed across prey species, consistent with a more generalist feeding pattern under comparatively heterogeneous prey availability. Rather than a strong dominance by a single prey type, IRI values were distributed more uniformly among species, suggesting reduced dependence on any single resource. Similar spatial shifts in prey composition, driven by relative local availability, have been documented in European Shag populations along the Norwegian coast (Barrett, 1991). Differences in local oceanographic conditions likely contribute to these spatial patterns. The upwelling system near Berlengas, influenced by Cabo Carvoeiro and the Nazaré Canyon, may enhance primary and secondary production, generating trophic hotspots characterised by abundant and predictable prey, promoting high foraging site fidelity and efficiency (O’Reilly et al., 2022; Morgan et al., 2019). In contrast, coastal areas, such as Arrábida, are subject to greater environmental variability and anthropogenic pressures, potentially leading to more heterogeneous prey distributions. Overall, these findings indicate that spatial variation in diet composition reflects local prey field structure and environmental context, reinforcing the species’ capacity to adjust foraging strategy to prevailing conditions. In addition to spatial differences, diet composition varied between breeding and non-breeding periods. Although no significant difference in IRI was detected, lower mean IRI and reduced variability during the breeding period suggest a tendency toward dietary specialization on a limited number of key prey species. This pattern is consistent with previous observations of breeding-period reliance on demersal, energy-rich prey (Xirouchakis et al., 2017; Consolo et al., 2011), likely reflecting the energetic constraints imposed by central-place foraging.

### Influence of diet on the reproductive productivity

Reproductive productivity exhibited a declining trend between 2016 and 2024, indicating reduced breeding success and/or chick survival over time. While multiple factors may contribute to this pattern, diet composition emerged as a key explanatory variable. In particular, higher biomass of energy-rich prey, especially sandeels, was positively associated with reproductive output, whereas reduced dominance of this prey corresponded with lower productivity. As long-lived central-place foragers, seabirds are highly sensitive to fluctuations in prey availability and energetic quality. Reduced access to high-quality prey can increase foraging effort, constrain chick provisioning rates, and, in some cases, lead to reproductive skipping (Weimerskirch et al., 2003; Ponchon et al., 2014; Lorentsen et al., 2019). In this study, increased reliance on lower-energy or less predictable species, such as Horse Mackerel, Sand smelt (*Atherina presbyter*), Cuckoo Wrasse (*Labrus mixtus*), was associated with reduced reproductive performance. Similar patterns have been documented in other seabird populations experiencing dietary shifts from high-to lower-quality prey (Wanless et al., 2005; Hjernquist & Hjernquist, 2010). Climate-driven alterations in prey distribution and abundance may further intensify these effects. Warming waters have been associated with the redistribution of sandeels and other key forage fish (MacLeod et al., 2007; Beaugrand et al., 2008), potentially reducing the availability of energetically profitable prey during critical breeding periods. Collectively, these findings highlight that both prey availability and energetic quality are critical determinants of reproductive success, emphasizing the importance of bottom-up trophic dynamics in shaping population trajectories.

### Impacts of Climate Change: Key Considerations

Seabirds play a pivotal role in marine ecosystems, functioning as top predators, nutrient transporters, and sensitive environmental indicators. In Portugal, the European Shag holds particular ecological importance, yet reproductive productivity has declined in recent years, contributing to its vulnerable conservations’ status (Nascimento et al., 2021; Young et al., 2023). Our findings suggest that environmental variability, particularly sea surface temperature and indicators of primary and secondary production, significantly influence diet composition, underscoring the susceptibility of this species to climate-mediated trophic shifts. Climate change can affect seabirds through bottom-up processes that alter primary productivity and propagate through higher trophic levels (Howells et al., 2017; Young et al., 2023), as well as through temporal or spatial mismatches between predator and prey (“match-mismatch”), which can disrupt phenology and breeding success. The observed significant relationships between IRI and chlorophyll-a concentration and zooplankton provide evidence that trophic dynamics in this system are environmentally regulated. Furthermore, the observed decline in prey dominance with increasing sea surface temperature suggests that continued ocean warming may reduce the availability or predictability of energetically profitable prey. Globally, comparable climate-driven trophic effects have been documented. For example, colder winters in the North Sea enhanced primary productivity and improved kittiwake breeding success (Frederiksen et al., 2006, 2007), whereas warming along the Norwegian coast caused phenological mismatches between zooplankton and herring, reducing puffin reproductive output (Durant et al., 2003). Species with narrow dietary ranges are more vulnerable than generalists, highlighting the importance of understanding the intrinsic and extrinsic factors that shape species-specific responses to climate change to guide effective seabird conservation strategies.

### Limitations and future directions

Despite its important findings, some limitations should be acknowledged in this study. Sample sizes were uneven across sites and periods, with substantially more samples collected in the Berlengas than in Arrábida, and during the breeding period than during the non-breeding period. Reduced winter sampling in the Berlengas likely reflects limited access due to adverse weather. Future studies should include Arrábida and additional colonies to enable temporal and spatial comparisons, and the use of GPS telemetry could improve links between foraging behaviour, diet composition, and reproductive success (Nascimento et al., 2021). Pellet analysis also posed methodological challenges. Otoliths may degrade, fragment, or closely resemble those of other species, increasing the risk of misidentification. Small prey remains may be lost from pellets (Veen et al., 2003), and secondary consumption can complicate dietary interpretation (Johnson et al., 1997). Complementary approaches - such as vertebrae and jaw analysis, stable isotope analysis (δ^13^C/δ^15^N), and DNA barcoding - can enhance dietary assessment and reduce potential errors (Barrett et al., 2007; Quigley et al., 2023). The Index of Relative Importance (IRI), while widely used, has inherent limitations due to redundancy among its components, which can bias prey importance rankings (MacDonald & Green, 1983; Moreno-Amich, 1996). Researcher-related limitations include restricted access to some nests, differences in methods among researchers, and missing biomass data in 2022 - 2023. Despite these constraints, the study provides valuable insights into the diet and productivity of European Shags in Portugal, offering a solid foundation for future long-term and comparative research.

## Conclusion

This study demonstrates that European Shag diet is primarily driven by environmental variability rather than temporal trends, with bottom-up processes influencing both prey composition and reproductive performance. Spatial and seasonal differences reflected local prey field structure and life-history constraints, while shifts away from energetically profitable prey were associated with reduced productivity. These findings highlight the sensitivity of coastal seabirds to climate-mediated trophic changes and underscore the importance of understanding prey dynamics for conservation in the face of ongoing environmental change.

## Supporting information

Figure S1 (Online Resource 1); Table S1, Online Resource 1;Table S2, Online Resource 1; Table S3, Online Resource 1

## Acknowledgements

Long-term support for the data collection and the processing of regurgitated pellets was provided by SPEA (Sociedade Portuguesa para o Estudo das Aves) and its volunteers. We thank the Instituto da Conservação da Natureza e das Florestas (ICNF) and the Nature Rangers of the Berlengas Natural Reserve and Arrábida Natural Park for providing logistical support, data collection, transportation, and permitting. Additional acknowledgement is extended to the Peniche Harbour Authority and the lighthouse keepers for transportation to Berlenga Grande Island and logistical assistance. Much of this work was carried out at the Laboratory of Biodiversity, Genetics and Evolution, Faculty of Sciences, University of Porto, which provided laboratory facilities and institutional support.

## References

Abookire AA, Piatt JF (2005) Oceanographic conditions structure forage fishes into lipid-rich and lipid-poor communities in lower Cook Inlet, Alaska, USA. Marine Ecology Progress Series 287:229–240. 10.3354/meps287229

Aebischer NJ (1995) Philopatry and colony fidelity of shags Phalacrocorax aristotelis on the east coast of Britain. Ibis 137:11–18. 10.1111/j.1474-919X.1995.tb03214.x

Al-Ismail S, McMinn M, Tuset V, Lombarte A, Alcover J (2013) Summer diet of European shags Phalacrocorax aristotelis desmarestii in southern Mallorca. Seabird 26:8–23. 10.61350/sbj.26.8

Almeida J, Godinho C, Leitão D, Lopes RJ (2022) Lista Vermelha das Aves de Portugal Continental. SPEA, ICNF, LabOR/UÉ, CIBIO/BIOPOLIS, Portugal

Ammar Y, Puntila-Dodd R, Tomczak MT, Nyström M, Blenckner T (2025) Novelty, variability, and resilience: Exploring adaptive cycles in a marine ecosystem under pressure. Ambio 54:1885–1901. 10.1007/s13280-025-02181-1

Aznar R, Sotillo MG, Cailleau S, Lorente P, Levier P, Amo-Baladrón A, Reffray G, Alvarez Fanjul E (2016) Strengths and weaknesses of the CMEMS forecasted and reanalysed solutions for the Iberia-Biscay-Ireland (IBI) waters. Journal of Marine Systems 159:1–14. 10.1016/j.jmarsys.2016.02.007

Banks DL, Fienberg SE (2003) Statistics, multivariate. In: Encyclopedia of Physical Science and Technology, 3rd edn. Elsevier, pp 851–889. 10.1016/B0-12-227410-5/00731-6

Barati, A., & Behrouzi-Rad, B. (2010). Breeding success of the great cormorant, Phalacrocorax carbo Linnaeus, 1758, at ramsar, northern iran: (Aves: Phalacrocoracidae). Zoology in the Middle East, 50(1), 41–46. 10.1080/09397140.2010.10638410

Barrett R, Rov N, Loen J, Montevecchi W (1990) Diets of shags Phalacrocorax aristotelis and cormorants P. carbo in Norway and possible implications for gadoid stock recruitment. Marine Ecology Progress Series 66:205–218. 10.3354/meps066205

Barrett RT (1991) Shags (Phalacrocorax aristotelis L.) as potential samplers of juvenile saithe (Pollachius virens (L.)) stocks in Northern Norway. Sarsia 76:153–156. 10.1080/00364827.1991.10413470

Barrett RT, Camphuysen KCJ, Anker-Nilssen T, Chardine JW, Furness RW, Garthe S, Hüppop O, Leopold MF, Montevecchi WA, Veit RR (2007) Diet studies of seabirds: A review and recommendations. ICES Journal of Marine Science 64:1675–1691. 10.1093/icesjms/fsm152

Beaugrand G, Edwards M, Brander K, Luczak C, Ibanez F (2008) Causes and projections of abrupt climate-driven ecosystem shifts in the North Atlantic. Ecology Letters 11:1157–1168. 10.1111/j.1461-0248.2008.01218.x

BirdLife International (2025) Site factsheet: Berlengas. Available at: https://datazone.birdlife.org/site/factsheet/berlengas

Consolo M, Privileggi N, Cimador B, Sponza S (2011) Dietary changes of Mediterranean Shags Phalacrocorax aristotelis desmarestii between the breeding and post-breeding seasons in the upper Adriatic Sea. Bird Study 58:461–472. 10.1080/00063657.2011.603290

Corona LS (2021) Shifts in the European shag diet along the Portuguese coast. MSc dissertation, University of Algarve, Portugal

Durant JM, Anker-Nilssen T, Stenseth NC (2003) Trophic interactions under climate fluctuations: The Atlantic puffin as an example. Proceedings of the Royal Society B: Biological Sciences 270:1461– 1466. 10.1098/rspb.2003.2397

Espinosa LA, Portela MM (2025) Red-hot Portugal: Mapping the increasing severity of exceptional maximum temperature events (1980–2024). Atmosphere 16:514. 10.3390/atmos16050514

Fernández N, Román J, Delibes M (2016) Variability in primary productivity determines metapopulation dynamics. Proceedings of the Royal Society B: Biological Sciences 283:20152998. 10.1098/rspb.2015.2998

Ferrari C, Evangelista C, Basiricò L, Castellani S, Biffani S, Bernabucci U (2025) Application of a generalized additive mixed model in time series study of dairy cow behavior under hot summer conditions. Journal of Dairy Science 108:1554–1572. 10.3168/jds.2024-25001

Franks PJS (1992) Phytoplankton blooms at fronts: Patterns, scales, and physical forcing mechanisms. Reviews in Aquatic Sciences 6:121–137.

Frederiksen M, Edwards M, Mavor RA, Wanless S (2007) Regional and annual variation in black-legged kittiwake breeding productivity is related to sea surface temperature. Marine Ecology Progress Series 350:137–143. 10.3354/meps07126

Frederiksen M, Edwards M, Richardson AJ, Halliday NC, Wanless S (2006) From plankton to top predators: Bottom-up control of a marine food web across four trophic levels. Journal of Animal Ecology 75:1259–1268. 10.1111/j.1365-2656.2006.01148.x

Froese R, Pauly D (eds) (2025) FishBase. World Wide Web electronic publication. Available at: https://www.fishbase.org

Furlan E, Stoklosa J, Griffiths J, Gust N, Ellis R, Huggins RM, Weeks AR (2012) Small population size and extremely low levels of genetic diversity in island populations of the platypus Ornithorhynchus anatinus. Ecology and Evolution 2:844–857. 10.1002/ece3.195

Furness RW, Tasker ML (1999) Diets of seabirds and consequences of changes in food supply. ICES Cooperative Research Report 232. 10.17895/ices.pub.5363

Gagnon K, Rothäusler E, Syrjänen A, Yli-Renko M, Jormalainen V (2013) Seabird guano fertilizes Baltic Sea littoral food webs. PLoS ONE 8:e61284. 10.1371/journal.pone.0061284

Gladstone-Gallagher RV, Thrush SF, Low JML, Pilditch CA, Ellis JI, Hewitt JE (2023) Toward a network perspective in coastal ecosystem management. Journal of Environmental Management 343:119007. 10.1016/j.jenvman.2023.119007

Grémillet D, Chauvin C, Wilson RP, Le Maho Y, Wanless S (2005) Unusual feather structure allows partial plumage wettability in diving great cormorants Phalacrocorax carbo. Journal of Avian Biology 36:57–63. 10.1111/j.0908-8857.2005.03331.x

Harris MP, Wanless S (1993) The diet of shags Phalacrocorax aristotelis during the chick-rearing period assessed by three methods. Bird Study 40:135–139. 10.1080/00063659309477138

Harris MP, Wanless S (1997) Breeding success, diet, and brood neglect in the kittiwake (Rissa tridactyla) over an 11-year period. ICES Journal of Marine Science 54:615–623. 10.1006/jmsc.1997.0241

Hart RK, Calver MC, Dickman CR (2002) The index of relative importance: An alternative approach to reducing bias in descriptive studies of animal diets. Wildlife Research 29:415–421. 10.1071/WR02009

Heath MR, Speirs DC, Steele JH (2014) Understanding patterns and processes in models of trophic cascades. Ecology Letters 17:101–114. 10.1111/ele.12200

Hillersøy G, Lorentsen SH (2012) Annual variation in the diet of breeding European shag (Phalacrocorax aristotelis) in central Norway. Waterbirds 35:420–429. 10.1675/063.035.0306

Hjernquist B, Hjernquist MB (2010) The effects of quantity and quality of prey on population fluctuations in three seabird species. Bird Study 57(1):19–25. 10.1080/00063650903029516

Howells RJ, Burthe SJ, Green JA, Harris MP, Newell MA, Butler A, Johns DG, Carnell EJ, Wanless S, Daunt F (2017) From days to decades: Short-and long-term variation in environmental conditions affect offspring diet composition of a marine top predator. Marine Ecology Progress Series 583:227–242. 10.3354/meps12343

Howells RJ, Burthe SJ, Green JA, Harris MP, Newell MA, Butler A, Wanless S, Daunt F (2018) Pronounced long-term trends in year-round diet composition of the European shag Phalacrocorax aristotelis. Marine Biology 165:193. 10.1007/s00227-018-3433-9

Jean-Michel L, Eric G, Romain B-B, Gilles G, Angélique M, Marie D, Clément B, Mathieu H, Olivier LG, Charly R, Tony C, Charles-Emmanuel T, Florent G, Giovanni R, Mounir B, Yann D, Pierre-Yves LT (2021) The Copernicus Global 1/12° oceanic and sea ice GLORYS12 reanalysis. Frontiers in Earth Science 9:698876. 10.3389/feart.2021.698876

Johnson, J. H., Ross, R. M., & Smith, D. R. (1997). Evidence of secondary consumption of invertebrate prey by Double-crested Cormorants. Waterbirds, 20(3), 547–551. 10.2307/1521608

Jones NM, McChesney GJ, Parker MW, Yee JL, Carter HR, Golightly RT (2008) Breeding phenology and reproductive success of the Brandt’s cormorant at three nearshore colonies in central California, 1997–2001. Waterbirds 31:505–514. 10.1675/1524-4695-31.4.505

Kennedy M, Spencer HG (2014) Classification of the cormorants of the world. Molecular Phylogenetics and Evolution 79:249–257. 10.1016/j.ympev.2014.06.020

Lomartire S, Marques JC, Gonçalves AMM (2021) The key role of zooplankton in ecosystem services: a perspective of interaction between zooplankton and fish recruitment. Ecol Indic 129:107867. 10.1016/j.ecolind.2021.107867

Lorentsen SH, Mattisson J, Christensen-Dalsgaard S (2019) Reproductive success in the European shag is linked to annual variation in diet and foraging trip metrics. Marine Ecology Progress Series 619:137–147. 10.3354/meps12949

Macdonald JS, Green RH (1983) Redundancy of variables used to describe importance of prey species in fish diets (Bay of Fundy). Canadian Journal of Fisheries and Aquatic Sciences 40:635–637.

MacLeod CD, Santos MB, Reid RJ, Scott BE, Pierce GJ (2007) Linking sandeel consumption and the likelihood of starvation in harbour porpoises in the Scottish North Sea: Could climate change mean more starving porpoises? Biology Letters 3:185–188. 10.1098/rsbl.2006.0588

Mann HB, Whitney DR (1947) On a test of whether one of two random variables is stochastically larger than the other. Annals of Mathematical Statistics 18:50–60. 10.1214/aoms/1177730491

Mitchell A, McCarthy E, Verspoor E (1998) Discrimination of the North Atlantic lesser sandeels Ammodytes marinus, A. tobianus, A. dubius and Gymnammodytes semisquamatus by mitochondrial DNA restriction fragment patterns. Fisheries Research 36:61–65. 10.1016/S0165-7836(98)00081-2

Morat F, Mante A, Drunat E, Dabat J, Bonhomme P, Harmelin-Vivien M, Letourneur Y (2014) Diet of Mediterranean European shag Phalacrocorax aristotelis desmarestii in a northwestern Mediterranean area: A competitor for local fisheries? Scientific Reports of the Port-Cros National Park 28:113–132.

Moreno-Amich R (1996) Feeding habits of longfin gurnard, Aspitrigla obscura (L. 1764), along the Catalan coast (northwestern Mediterranean). Hydrobiologia 324:219–228.

Morgan EA, Hassall C, Redfern CPF, Bevan RM, Hamer KC (2019) Individuality of foraging behaviour in a short-ranging benthic marine predator: Incidence and implications. Marine Ecology Progress Series 609:209–219. 10.3354/meps12819

Nascimento T, Oliveira N, Luís A (2021) Hey, that’s my fish - overlap in prey composition between European shag and local fisheries in Portugal. Ardea 109:77–90. 10.5253/arde.v109i1.a10

Nascimento T, Oliveira N, Luís A (2023) Spatial overlap between the European shag and commercial fisheries in a special protected area: Implications for conservation. Fisheries Research 263:106689. 10.1016/j.fishres.2023.106689

Nascimento TSL (2018) O papel da pesca comercial na conservação da população de galheta Phalacrocorax aristotelis do arquipélago das Berlengas. MSc dissertation, University of Aveiro, Portugal.

O’Reilly L, Fentimen R, Butschek F, Titschack J, Lim A, Moore N, O’Connor OJ, Appah J, Harris K, Vennemann T, Wheeler AJ (2022) Environmental forcing by submarine canyons: Evidence between two closely situated cold-water coral mounds (Porcupine Bank Canyon and Western Porcupine Bank, NE Atlantic). Marine Geology 454:106930. 10.1016/j.margeo.2022.106930

Pallavi P, Parthasarathy D, Narayanan K, Inamdar AB, Budakoti S (2024) Examining the principal factors that limit chlorophyll-a concentration across coastal waters of northern Maharashtra state using a robust generalised additive model. 10.1016/j.rsma.2024.103693

Perkins A, Ratcliffe N, Suddaby D, Ribbands B, Smith C, Ellis P, Bolton M (2018) Combined bottom-up and top-down pressures drive catastrophic population declines of Arctic skuas in Scotland. Journal of Animal Ecology 87:1573–1586. 10.1111/1365-2656.12890

Piatt JF, Sydeman WJ, Wiese F (2007) Introduction: A modern role for seabirds as indicators. Marine Ecology Progress Series 352:199–204. 10.3354/meps07070

Pinkas L, Oliphant MS, Iverson ILK (1971) Food habits of albacore, bluefin tuna and bonito in Californian waters. California Department of Fish and Game Fish Bulletin 152:1–105.

Ponchon A, Grémillet D, Christensen-Dalsgaard S, Erikstad KE, Barrett RT, Reiertsen TK, McCoy KD, Tveraa T, Boulinier T (2014) When things go wrong: Intra-season dynamics of breeding failure in a seabird. Ecosphere 5(1). 10.1890/ES13-00233.1

Provencher JF, Borrelle SB, Bond AL, Lavers JL, Franeker JA, Kühn S, Hammer S, Avery-Gomm S, Mallory ML (2019) Recommended best practices for plastic and litter ingestion studies in marine birds: Collection, processing, and reporting. FACETS 4:111–148. 10.1139/facets-2018-0043

Quigley LA, Caiger PE, Govindarajan AF, McMonagle H, Jech JM, Lavery AC, Llopiz JK (2023) Otolith characterization and integrative species identification of adult mesopelagic fishes from the western North Atlantic Ocean. Frontiers in Marine Science 10:1217779. 10.3389/fmars.2023.1217779

Rajpar MN, Ozdemir I, Zakaria M, Sheryar S, Rab A (2018) Seabirds as bioindicators of marine ecosystems. In: Seabirds. InTech. 10.5772/intechopen.75458

Ratnarajah L, Abu-Alhaija R, Atkinson A, Batten S, Bax NJ, Bernard KS, Canonico G, Cornils A, Everett JD, Grigoratou M, Ishak NHA, Johns D, Lombard F, Muxagata E, Ostle C, Pitois S, Richardson AJ, Schmidt K, Stemmann L, Swadling KM, Yang G, Yebra L (2023) Monitoring and modelling marine zooplankton in a changing climate. Nat Commun 14:2602. 10.1038/s41467-023-36241-5

Sandvik H (2004) Life-history and breeding biology of seabirds in a changing environment: A comparative approach. PhD thesis, Norwegian University of Science and Technology, Tromsø

Sherley RB, Ludynia K, Underhill LG, Jones R, Kemper J (2012) Storms and heat limit the nest success of Bank cormorants: Implications of future climate change for a surface-nesting seabird in southern Africa. Journal of Ornithology 153:441–455. 10.1007/s10336-011-0760-8

Simeoni C, Furlan E, Pham HV, Critto A, de Juan S, Trégarot E, Cornet CC, Meesters E, Fonseca C, Botelho AZ, Krause T, N’Guetta A, Cordova FE, Failler P, Marcomini A (2023) Evaluating the combined effect of climate and anthropogenic stressors on marine coastal ecosystems: Insights from a systematic review of cumulative impact assessment approaches. Science of the Total Environment 861: 160687 10.1016/j.scitotenv.2022.160687

Staudinger MD, Goyert H, Suca JJ, Coleman K, Welch L, Llopiz JK, Wiley D, Altman I, Applegate A, Auster P, Baumann H, Beaty J, Boelke D, Kaufman L, Loring P, Moxley J, Paton S, Powers K, Richardson D, Robbins J, Runge J, Smith B, Spiegel C, Steinmetz H (2020) The role of sand lances (Ammodytes sp.) in the Northwest Atlantic ecosystem: A synthesis of current knowledge with implications for conservation and management. Fish and Fisheries 21:522–556. 10.1111/faf.12445

Ullah H, Nagelkerken I, Goldenberg SU, Fordham DA (2018) Climate change could drive marine food web collapse through altered trophic flows and cyanobacterial proliferation. PLoS Biology 16: e2003446 10.1371/journal.pbio.2003446

van der Kooij J, Scott BE, Mackinson S (2008) The effects of environmental factors on daytime sandeel distribution and abundance on the Dogger Bank. J Sea Res 60:201–209. 10.1016/j.seares.2008.07.003

Van Guilder MA, Seefelt NE (2013) Double-crested cormorant (Phalacrocorax auritus) chick bioenergetics following round goby (Neogobius melanostomus) invasion and implementation of cormorant population control. Journal of Great Lakes Research 39:153–161. 10.1016/j.jglr.2012.12.019

Veen J, Peeters J, Leopold MF, van Damme CJG, Veen T (2003) Les oiseaux piscivores comme indicateurs de la qualité de l’environnement marin: Suivi des effets de la pêche littorale en Afrique du Nord-Ouest. Alterra Report 666, Alterra.

Velando A, Freire J (1999) Intercolony and seasonal differences in the breeding diet of European shags Phalacrocorax aristotelis on the Galician coast (NW Spain). Marine Ecology Progress Series 188:225–236. 10.3354/meps188225

Vereshchaka A (2024) Navigating the zooplankton realm: Oceans of diversity beneath the sea surface. Diversity 16:717. 10.3390/d16120717

Wanless, S., Harris, M. P., Redman, P., & Speakman, J. R. (2005). Low energy values of fish as a probable cause of a major seabird breeding failure in the North Sea. Marine Ecology Progress Series, 294, 1–8. 10.3354/meps294001

Weeks SJ, Steinberg C, Congdon BC (2013) Oceanography and seabird foraging: Within-season impacts of increasing sea-surface temperature on the Great Barrier Reef. Marine Ecology Progress Series 490:247–254. 10.3354/meps10398

Weimerskirch H, Ancel A, Caloin M, Zahariev A, Spagiari J, Kersten M, et al. (2003) Foraging efficiency and adjustment of energy expenditure in a pelagic seabird provisioning its chick. Journal of Animal Ecology 72:500–508.

Winslade P (1974) Behavioural studies on the lesser sandeel Ammodytes marinus (Raitt) I. The effect of food availability on activity and the role of olfaction in food detection. Journal of Fish Biology 6:565–576. 10.1111/j.1095-8649.1974.tb05100.x

Xirouchakis SM, Kasapidis P, Christidis A, Andreou G, Kontogeorgos I, Lymberakis P (2017) Status and diet of the European shag (Mediterranean subspecies) Phalacrocorax aristotelis desmarestii in the Libyan Sea (south Crete) during the breeding season. Marine Ornithology 45:1–9. 10.5038/2074-1235.45.1.1192

Young LC, VanderWerf E (2022) Conservation of marine birds. Elsevier. 10.1016/C2020-0-03628-5

