## Supplementary material for "Diet and breeding productivity in European Shag (*Gulosus aristotelis*): insights from two Portuguese colonies": Figure S1 (Online Resource 1); Table S1, Online Resource 1;Table S2, Online Resource 1; Table S3, Online Resource 1

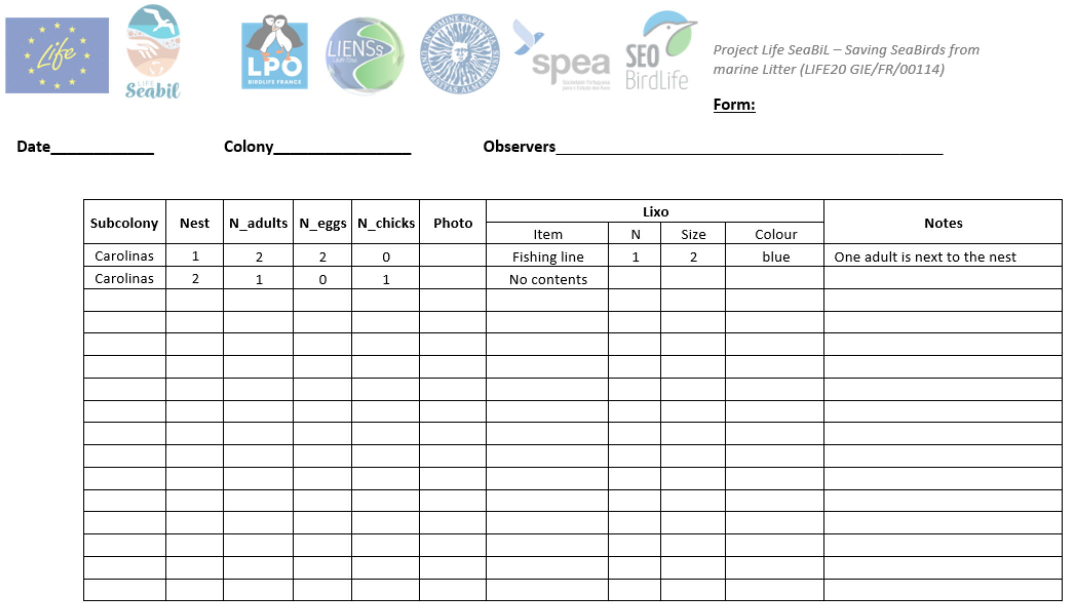

Project Life SeaBiL – Saving SeaBirds from marine Litter (LIFE20 GIE/FR/00114)

**Form:**

**Date**___________ **Colony**______________ **Observers**________________________________________

| Subcolony | Nest | N_adults | N_eggs | N_chicks | Photo | Lixo | | | | Notes |
|---|---|---|---|---|---|---|---|---|---|---|
| | | | | | | Item | N | Size | Colour | |
| Carolinas | 1 | 2 | 2 | 0 | | Fishing line | 1 | 2 | blue | One adult is next to the nest |
| Carolinas | 2 | 1 | 0 | 1 | | No contents | | | | |

*Figure S1. Monitoring sheets were used to record the number of adults, eggs, hatched chicks, and live juveniles to assess the productivity of the European Shag (Gulosus aristotelis).*

*Table S 1. Equations for Length-weight relationship regression based on otolith's length, TL = fish total length; OL = otolith length; TW= fish total weight*

| Family/Species | Length Equation (cm) - FL | Biomass Equation (g) - FW | Reference |
|---|---|---|---|
| Ammodytidae | $\frac{(OL-0,510)}{0,141}$ | $0,00660 \text{ x } (FL^{2,697})$ | (Wilson et al., 2017; Froese & Pauly, 2018) |
| Argentinidae<br>*Argentina sphyraena* | $\frac{(40,72 \text{ x } (OL^{0,83}))}{10}$ | $0,0083 \text{ x } (FL^{3,266})$ | (Giménez et al., 2016; Torres et al., 2012) |
| Atherinidae<br>*Atherina boyeri*<br>*Atherina presbyter* | $\frac{(52,11 \text{ x } (OL^{0,38}))}{10}$ | $0,0033 \text{ x } (FL^{3,35})$<br>$0,0055 \text{ x } (FL^{3,09})$ | (Giménez et al., 2016; Pombo et al., 2005) |
| Blenniidae<br>*Parablennius sp.*<br>*Parablennius pilicornis* | $\frac{(41,85 \text{ x } (OL^{1,19}))}{10}$<br>$\frac{(44,31 \text{ x } (OL^{1,02}))}{10}$ | $0,01223 \text{ x } (FL^{2,7693})$<br>$0,01223 \text{ x } (FL^{2,7693})$ | (Giménez et al., 2016; Valle et al., 2002) |
| Bothidae<br>*Arnoglossus imperialis* | $\frac{(51,64 \text{ x } (OL^{0,92}))}{10}$ | $0,0074 \text{ x } (FL^{3,024})$ | (Giménez et al., 2016; Torres et al., 2012) |
| Engraulidae<br>*Engraulis encrasicolus* | $\frac{(46,87 \text{ x } (OL^{0,92}))}{10}$ | $0,0049 \text{ x } (FL^{3,125})$ | (Giménez et al., 2016; Torres et al., 2012) |
| Gadidae<br>*Trisopterus luscus*<br>*Trisopterus minutus* | $\frac{(33,73 \text{ x } (OL - 94,54}{10}$ [illegible] | $0,00000433 \text{ x } (FL \text{ x } 10)$ [illegible]<br>$0,0042 \text{ x } (FL^{3,3425})$ | (Granadeiro and Silva, 2000; Wilson et al., 2017; Morey et al. 2003) |

| Family/Species | Length Equation (cm) - FL | Biomass Equation (g) - FW | Reference |
|---|---|---|---|
| | $\frac{OL - 1,718}{0,362}$ | | |
| Gobidae<br>*Gobius paganellus*<br>*Gobius sp.* | $2,296 \text{ x } OL + 0,138$ | $0,00660 \text{ x } (FL^{3,5738})$<br>$0,0089 \text{ x } (FL^{3,0853})$ | (İlkyaz et al., 2011; Morey et al. 2003) |
| Triglidae<br>*Trigla lyra* | $\frac{(20,78 \text{ x } (OL^{1,66}))}{10}$ | $0,0092 \text{ x } (FL^{2,9308})$ | (Giménez et al., 2016; Mendes et al. 2004) |
| Trachinidae<br>*Trachinus draco*<br>*Echiichtys vipera* | $\frac{(15,66 \text{ x } (OL^{1,25}))}{10}$<br>$\frac{(16,43 \text{ x } (OL^{1,22}))}{10}$ | $0,0042 \text{ x } (FL^{3,119})$<br>$0,01050 \text{ x } (FL^{3,05})$ | (Giménez et al., 2016; Santos et al., 2002; Viegas et al., 2009) |
| Carangidae<br>*Trachurus trachurus* | $\frac{(34,01 \text{ x } OL - 28,9}{10}$ | $0,00001,77 \text{ x } (FL \text{ x } 10)^{2}$ | (Granadeiro and Silva, 2000) |
| Clupeidae<br>*Sardina pilchardus* | $\frac{(55,75 \text{ x } OL^{0,95})}{10}$ | $0,00000692 \text{ x } (FL \text{ x } 10)$ | (Giménez et al., 2016; Granadeiro and Silva, 2000) |
| Soleidae<br>*Pegusa lascaris*<br>*Microchirus sp* | $\frac{(58,12 \text{ x } OL^{1,11})}{10}$<br>$\frac{(44,56 \text{ x } OL^{0,97})}{10}$ | $0,0070 \text{ x } FL^{3,13}$<br>$0,0026 \text{ x } FL^{3,285}$ | (Giménez et al., 2016; Mendes et al. 2004) |
| Labridae<br>*Coris julis* | $\frac{(31,63 \text{ x } OL^{1,56})}{10}$ | $0,00001409 \text{ x } FLx\ 10^{2,9}$ | (Giménez et al., 2016; Gonçalves et al., 1997) |
| *Ctenolabrus rupestris* | $\frac{(31,63 \text{ x } OL^{1,56})}{10}$ | $0,00001409 \text{ x } FLx\ 10^{2,9}$ | (Giménez et al., 2016; Gonçalves et al., 1997) |
| *Labrus bergylta* | $\frac{(67,97 \text{ x } OL^{31,24})}{10}$ | $0,0141 \text{ x } FL^{3,039}$ | (Härkönen, 1986; Morato et al., 2001) |
| *Labrus merula* | $\frac{(57,52 \text{ x } OL - 4,76}{10}$ | $0,0076 \text{ x } FL^{3,1862}$ | (Giménez et al., 2016; Morey et al. 2003) |
| *Labrus mixtus* | $\frac{(52,12 \text{ x } OL^{4,76})}{10}$ | $0,0050 \text{ x } FL^{3,254}$ | (Härkönen, 1986; Mendes et al. 2004) |
| *Symphodus rossali* | $4,3583 \text{ x } OL^{0,9869}$ | $0,0155 \text{ x } FL^{3,017}$ | (Altin and Ayyildiz 2017; Santos et al., 2002) |
| *Symphodus cinereus* | $\frac{(48,27 \text{ x } OL^{0,94})}{10}$ | $0,0075 \text{ x } FL^{3,214}$ | (Giménez et al., 2016; Morey et al. 2003) |
| *Symphodus mediterraneus* | $\frac{(31,25 \text{ x } (OL^{1,47})}{10}$ | $0,02121 \text{ x } FL^{2,8798}$ | (Giménez et al., 2016; Valle et al.,2002) |
| *Symphodus melops* | $\frac{(48,27 \text{ x } (OL^{0,94})}{10}$ | $0,01120 \text{ x } FL^{3,17}$ | (Altin and Ayyildiz, 2017; Froese and Pauly, 2018) |
| *Symphodus sp.* | $\frac{(31,25 \text{ x } OL^{1,47})}{10}$ | $0,0155 \text{ x } FL^{3,017}$ | (Giménez et al., 2016; Santos et al., 2002) |
| Phycidae<br>*Phycis sp.* | $\frac{(1,89 \text{ x } OL^{2,01})}{10}$ | $0,0064 \text{ x } FL^{3,149}$ | (Mendes et al., 2004) |
| *Gaidropsarus mediterraneus* | $\frac{(36,91 \text{ x } OL^{1,05})}{10}$ | $0,0012 \text{ x } FL^{3,616}$ | (Giménez et al., 2016; Kasapoglu et al., 2014) |

| Family/Species | Length Equation (cm) - FL | Biomass Equation (g) - FW | Reference |
|---|---|---|---|
| *Ciliata mustela* | $\frac{(92,29 \times OL - 74,6}{10}$ | $1,0736 \times FL^{3,444}$ | (Härkönen 1986) |
| Serranidae | | | |
| *Serranus hepatus* | $\frac{(16,99 \times OL^{1,15})}{10}$ | $0,0110 \times FL^{3,190}$ | (Giménez et al., 2016; Santos et al., 2002 |
| *Serranus cabrilla* | $\frac{(21,75 \times OL^{1,11})}{10}$ | $0,0092 \times FL^{3,0658}$ | (Morey et al. 2003) |
| Sparidae | | | |
| *Diplodus sp.* | $\frac{(20,04 \times OL^{1,27})}{10}$ | $0,0132 \times FL^{3,096}$ | (Giménez et al., 2016; Santos et al., 2002) |
| *Diplodus annularis* | $\frac{(20,04 \times OL^{1,27})}{10}$ | $0,0132 \times FL^{3,096}$ | (Giménez et al., 2016; Santos et al., 2002) |
| *Diplodus vulgaris* | $\frac{(17,18 \times OL^{1,13})}{10}$ | $0,00001756 \times (FLx10)^{2,}$ | (Giménez et al., 2016; Gonçalves et al., 1997) |
| *Spondyliosoma cantharus* | $\frac{(20,21 \times OL^{1,16})}{10}$ | $0,0158 \times FL^{2,9957}$ | (Giménez et al., 2016; Morey et al. 2003) |
| *Boops boops* | $\frac{(23,9 \times OL^{1,16})}{10}$ | $0,00000758 \times (FLx10)^{3,}$ | (Giménez et al., 2016; (Gonçalves et al. 1997) |
| *Dentex sp.* | $\frac{(4,59 \times OL^{1,82})}{10}$ | $0,0113 \times FL^{3,0349}$ | (Giménez et al., 2016; Morey et al. 2003) |
| *Diplodus sargus* | $\frac{(20,25 \times OL^{1,28})}{10}$ | $0,00001423 \times (FLx10)^{3,}$ | (Giménez et al., 2016; (Gonçalves et al. 1997) |
| *Sparus aurata* | $\frac{(18,04 \times OL^{1,32})}{10}$ | $0,00001827 \times FL^{2,96}$ | (Giménez et al., 2016; (Gonçalves et al. 1997) |
| Scorpaenidae | | | |
| *Scorpaena elongata* | $\frac{(12,6x\ OL^{1,2})}{10}$ | $0,0148 \times FL^{3,017}$ | (Giménez et al., 2016; Meiners-Mandujano et al., 2018) |

*References:*

Altin, A., & Ayyildiz, H. (2017). Relationships between total length and otolith measurements for 36 fish species from Gökçeada Island, Turkey. Journal of Applied Ichthyology, 1-6. https://doi.org/10.1111/jai.13509

Froese, R., & Pauly, D. (2018). FishBase. Retrieved from http://www.fishbase.org

Giménez, J., Manjabacas, A., Tuset, V. M., & Lombarte, A. (2016). Relationships between otolith and fish size from Mediterranean and north-eastern Atlantic species to be used in predator-prey studies. Journal of Fish Biology, 89, 2195-2202. https://doi.org/10.1111/jfb.13115

Gonçalves, J. M. S., Bentes, L., Lino, P. G., Ribeiro, J., Canário, A. V. M., & Erzini, K. (1997). Weight-length relationships for selected fish species of the small-scale demersal fisheries of the south and south-west coast of Portugal. *Fisheries Research, 30*, 253-256. https://doi.org/10.1016/S0165-7836(96)00569-3

Granadeiro, J. P., & Silva, M. A. (2000). The use of otoliths and vertebrae in the identification and size-estimation of fish in predator-prey studies. *Cybium, 24*, 383-393.

Härkönen, T. (1986). *Guide to the otoliths of the bony fishes of northeast Atlantic*. Danbiu ApS.

Meiners-Mandujano, C., Fernández-Peralta, L., Faraj, A., & García-Cancela, R. (2018). Length-weight relations of 15 deep-sea fish species (Actinopterygii) from the north-western African continental slope. *Acta Ichthyologica et Piscatoria, 48*(2), 195-198.

Mendes, B., Fonseca, P., & Campos, A. (2004). Weight-length relationships for 46 fish species of the Portuguese west coast. *Journal of Applied Ichthyology, 20*, 355-361. https://doi.org/10.1111/j.1439-0426.2004.00559.x

Morey, G., Moranta, J., Massutí, E., Grau, A., Linde, M., Riera, F., & Morales-Nin, B. (2003). Weight-length relationships of littoral to lower slope fishes from the western Mediterranean. *Fisheries Research, 62*, 89-96. https://doi.org/10.1016/S0165-7836(02)00250-3

Pombo, L., Elliot, M., & Rebelo, J. (2005). Ecology, age and growth of *Atherina boyeri* and *Atherina presbyter* in the Ria de Aveiro, Portugal. *Cybium, 29*, 47-55.

Santos, M. N., Gaspar, M. B., Vasconcelos, P., & Monteiro, C. C. (2002). Weight-length relationships for 50 selected fish species of the Algarve coast (southern Portugal). *Fisheries Research, 59*, 289-295. https://doi.org/10.1016/S0165-7836(01)00401-5

Veiga, P., Machado, D., Almeida, C., Bentes, L., Monteiro, P., Oliveira, F., ... Gonçalves, J. M. S. (2009). Weight-length relationships for 54 species of the Arade estuary, southern Portugal. *Journal of Applied Ichthyology, 25*(4), 493-496.

Wilson, L. J., Grellier, K., & Hammond, P. S. (2017). Improved estimates of digestion correction factors and passage rates for harbor seal (*Phoca vitulina*) prey in the northeast Atlantic. *Marine Mammal Science, 33*, 1149-1169. https://doi.org/10.1111/mms.12436

*Table S2. Identification of prey and its qualitative analysis (F.N = numerical frequency; F.O = frequency of occurrence; F.B = frequency of biomass) across reproductive and non-reproductive seasons in both Berlenga (2016–-2024) and Arrábida (2020-2021).*

| Year | Period | Area | Species | FN | FO | FB |
|---|---|---|---|---|---|---|
| 2016 | Breeding | Berlengas | *Trisopterus luscus* | 47.27 | 75 | 94.28 |
| 2016 | Breeding | Berlengas | Gadidae | 3.64 | 12.5 | 8.61 |
| 2016 | Breeding | Berlengas | Labridae | 16.36 | 37.5 | 36.45 |
| 2016 | Breeding | Berlengas | *Ciliata mustela* | 1.82 | 12.5 | 0.26 |
| 2016 | Breeding | Berlengas | Gaidropsarus sp. | 5.45 | 37.5 | 28.26 |
| 2016 | Breeding | Berlengas | *Labrus bergylta* | 1.82 | 12.5 | 2.60 |
| 2016 | Breeding | Berlengas | *Labrus mixtus* | 1.82 | 12.5 | 0.83 |

| Year | Period | Area | Species | FN | FO | FB |
|---|---|---|---|---|---|---|
| 2016 | Breeding | Berlengas | *Coris julis* | 3.64 | 12.5 | 0.58 |
| 2016 | Breeding | Berlengas | *Ammodytidae* | 3.64 | 25 | 3.21 |
| 2016 | Breeding | Berlengas | *Acantholabrus palloni* | 1.82 | 12.5 | 1.18 |
| 2016 | Breeding | Berlengas | *Symphodus bailloni* | 5.45 | 37.5 | 31.48 |
| 2016 | Breeding | Berlengas | *Boops boops* | 1.82 | 12.5 | 3.27 |
| 2016 | Breeding | Berlengas | *Symphodus melops* | 3.64 | 12.5 | 1.56 |
| 2016 | Non-Breeding | Berlengas | *Trisopterus luscus* | 4.17 | 20 | 8.42 |
| 2016 | Non-Breeding | Berlengas | *Coris julis* | 33.33 | 40 | 58.01 |
| 2016 | Non-Breeding | Berlengas | *Ammodytidae* | 4.17 | 20 | 0.09 |
| 2016 | Non-Breeding | Berlengas | *Gobius sp.* | 4.17 | 20 | 0.03 |
| 2016 | Non-Breeding | Berlengas | *Diplodus vulgaris* | 4.17 | 20 | 6.05 |
| 2016 | Non-Breeding | Berlengas | *Serranus cabrilla* | 8.33 | 40 | 57.53 |
| 2016 | Non-Breeding | Berlengas | *Ciliata mustela* | 8.33 | 40 | 36.99 |
| 2016 | Non-Breeding | Berlengas | *Gobiidae* | 12.50 | 20 | 0.41 |
| 2016 | Non-Breeding | Berlengas | *Centrolabrus trutta* | 8.33 | 20 | 7.46 |
| 2016 | Non-Breeding | Berlengas | *Spondyliosoma cantharus* | 4.17 | 20 | 25.60 |
| 2017 | Breeding | Berlengas | *Diplodus vulgaris* | 0.11 | 1.20 | 0.17 |
| 2017 | Breeding | Berlengas | *Gaidropsarus sp.* | 0.11 | 2.41 | 0.10 |
| 2017 | Breeding | Berlengas | *Labridae* | 0.28 | 4.82 | 0.31 |
| 2017 | Breeding | Berlengas | *Ammodytidae* | 73.21 | 66.27 | 58.62 |
| 2017 | Breeding | Berlengas | *Ciliata mustela* | 0.17 | 3.61 | 2.31 |
| 2017 | Breeding | Berlengas | *Labrus bergylta* | 0.45 | 7.23 | 2.30 |
| 2017 | Breeding | Berlengas | *Centrolabrus trutta* | 0.57 | 7.23 | 1.00 |
| 2017 | Breeding | Berlengas | *Coris julis* | 16.46 | 39.76 | 11.19 |
| 2017 | Breeding | Berlengas | *Argentina sphyraena* | 0.06 | 1.20 | 0.13 |
| 2017 | Breeding | Berlengas | *Sparidae* | 0.40 | 3.61 | 4.13 |
| 2017 | Breeding | Berlengas | *Gaidropsarus biscayensis* | 0.11 | 1.20 | 0.13 |
| 2017 | Breeding | Berlengas | *Gobius paganellus* | 0.11 | 2.41 | 0.02 |
| 2017 | Breeding | Berlengas | *Pagellus bogaraveo* | 0.06 | 1.20 | 0.02 |
| 2017 | Breeding | Berlengas | *Symphodus bailloni* | 1.31 | 8.43 | 2.05 |
| 2017 | Breeding | Berlengas | *Boops boops* | 0.45 | 8.43 | 0.84 |
| 2017 | Breeding | Berlengas | *Dentex sp.* | 0.17 | 2.41 | 0.03 |
| 2017 | Breeding | Berlengas | *Gaidropsarus mediterraneus* | 0.34 | 3.61 | 0.44 |
| 2017 | Breeding | Berlengas | *Sparus aurata* | 0.11 | 2.41 | 0.29 |
| 2017 | Breeding | Berlengas | *Symphodus melops* | 1.36 | 14.46 | 2.19 |
| 2017 | Breeding | Berlengas | *Trisopterus luscus* | 0.06 | 1.20 | 0.13 |
| 2017 | Breeding | Berlengas | *Arnoglossus imperialis* | 0.06 | 1.20 | 0.05 |
| 2017 | Breeding | Berlengas | *Serranus cabrilla* | 0.28 | 4.82 | 3.11 |
| 2017 | Breeding | Berlengas | *Sardina pilchardus* | 0.11 | 1.20 | 0.43 |
| 2017 | Breeding | Berlengas | *Atherina boyeri* | 0.23 | 1.20 | 0.12 |
| 2017 | Breeding | Berlengas | *Symphodus cinereus* | 0.06 | 1.20 | 0.03 |
| 2017 | Breeding | Berlengas | Atherina presbyter | 0.11 | 2.41 | 0.05 |
| 2017 | Breeding | Berlengas | *Labrus merula* | 0.74 | 4.82 | 3.14 |
| 2017 | Breeding | Berlengas | *Labrus mixtus* | 0.28 | 3.61 | 0.20 |
| 2017 | Breeding | Berlengas | *Engraulis encrasicolus* | 0.28 | 1.20 | 0.08 |
| 2017 | Breeding | Berlengas | *Diplodus sargus* | 0.17 | 2.41 | 1.33 |
| 2017 | Breeding | Berlengas | *Trigla lyra* | 0.06 | 1.20 | 0.19 |
| 2017 | Breeding | Berlengas | *Acantholabrus palloni* | 0.17 | 1.20 | 0.42 |
| 2017 | Breeding | Berlengas | *Gadidae* | 0.11 | 1.20 | 1.43 |

| Year | Period | Area | Species | FN | FO | FB |
|---|---|---|---|---|---|---|
| 2017 | Breeding | Berlengas | *Diplodus sp.* | 0.17 | 1.20 | 1.54 |
| 2017 | Breeding | Berlengas | *Triglidae* | 0.40 | 1.20 | 1.30 |
| 2017 | Breeding | Berlengas | *Trisopterus minutus* | 0.11 | 1.20 | 0.17 |
| 2017 | Non-Breeding | Berlengas | *Ammodytidae* | 43.53 | 34.78 | 7.79 |
| 2017 | Non-Breeding | Berlengas | *Argentina sphyraena* | 0.59 | 4.35 | 0.18 |
| 2017 | Non-Breeding | Berlengas | *Atherina boyeri* | 4.71 | 8.70 | 0.32 |
| 2017 | Non-Breeding | Berlengas | *Trachurus trachurus* | 0.59 | 4.35 | 5.21 |
| 2017 | Non-Breeding | Berlengas | *Trisopterus luscus* | 2.94 | 13.04 | 19.69 |
| 2017 | Non-Breeding | Berlengas | *Trisopterus minutus* | 1.76 | 8.70 | 12.78 |
| 2017 | Non-Breeding | Berlengas | *Gadidae* | 0.59 | 4.35 | 1.87 |
| 2017 | Non-Breeding | Berlengas | *Gobius paganellus* | 0.59 | 4.35 | 0.01 |
| 2017 | Non-Breeding | Berlengas | *Centrolabrus trutta* | 0.59 | 4.35 | 0.30 |
| 2017 | Non-Breeding | Berlengas | *Coris julis* | 18.24 | 34.78 | 3.21 |
| 2017 | Non-Breeding | Berlengas | *Labrus bergylta* | 0.59 | 4.35 | 0.40 |
| 2017 | Non-Breeding | Berlengas | *Labrus mixtus* | 1.76 | 8.70 | 1.15 |
| 2017 | Non-Breeding | Berlengas | *Symphodus bailloni* | 4.71 | 21.74 | 1.75 |
| 2017 | Non-Breeding | Berlengas | *Symphodus cinereus* | 1.76 | 4.35 | 0.10 |
| 2017 | Non-Breeding | Berlengas | *Symphodus melops* | 1.76 | 13.04 | 0.62 |
| 2017 | Non-Breeding | Berlengas | *Labridae* | 0.59 | 4.35 | 0.16 |
| 2017 | Non-Breeding | Berlengas | *Ciliata mustela* | 0.59 | 4.35 | 3.20 |
| 2017 | Non-Breeding | Berlengas | *Gaidropsarus biscayensis* | 0.59 | 4.35 | 0.17 |
| 2017 | Non-Breeding | Berlengas | *Serranus cabrilla* | 1.76 | 8.70 | 1.40 |
| 2017 | Non-Breeding | Berlengas | *Boops boops* | 1.18 | 8.70 | 8.98 |
| 2017 | Non-Breeding | Berlengas | *Diplodus sargus* | 0.59 | 4.35 | 11.14 |
| 2017 | Non-Breeding | Berlengas | *Diplodus vulgaris* | 4.12 | 8.70 | 2.78 |
| 2017 | Non-Breeding | Berlengas | *Spondyliosoma cantharus* | 0.12 | 8.70 | 16.61 |
| 2018 | Breeding | Berlengas | *Ammodytidae* | 2.33 | 11.11 | 0.16 |
| 2018 | Breeding | Berlengas | *Trachurus trachurus* | 6.98 | 22.22 | 67.85 |
| 2018 | Breeding | Berlengas | *Sardina pilchardus* | 4.65 | 22.22 | 4.92 |
| 2018 | Breeding | Berlengas | *Gaidropsarus biscayensis* | 2.33 | 11.11 | 0.97 |
| 2018 | Breeding | Berlengas | *Gaidropsarus sp.* | 4.65 | 11.11 | 3.50 |
| 2018 | Breeding | Berlengas | *Gobius sp.* | 2.33 | 11.11 | 0.20 |
| 2018 | Breeding | Berlengas | *Acantholabrus palloni* | 4.65 | 11.11 | 1.58 |
| 2018 | Breeding | Berlengas | *Coris julis* | 37.21 | 44.44 | 4.68 |
| 2018 | Breeding | Berlengas | *Labrus bergylta* | 2.33 | 11.11 | 2.52 |
| 2018 | Breeding | Berlengas | *Labrus mixtus* | 2.33 | 11.11 | 1.22 |
| 2018 | Breeding | Berlengas | *Symphodus bailloni* | 4.65 | 22.22 | 15.58 |
| 2018 | Breeding | Berlengas | *Symphodus melops* | 4.65 | 22.22 | 3.53 |
| 2018 | Breeding | Berlengas | *Ciliata mustela* | 2.33 | 11.11 | 3.68 |
| 2018 | Breeding | Berlengas | *Pegusa lascaris* | 2.33 | 11.11 | 3.40 |
| 2018 | Non-Breeding | Berlengas | *Trisopterus luscus* | 36.36 | 100 | 56.19 |
| 2018 | Non-Breeding | Berlengas | *Coris julis* | 18.18 | 50 | 2.24 |
| 2018 | Non-Breeding | Berlengas | *Gaidropsarus mediterraneus* | 36.36 | 100 | 31.58 |
| 2018 | Non-Breeding | Berlengas | *Serranus cabrilla* | 9.09 | 50 | 9.99 |
| 2019 | Breeding | Berlengas | *Ammodytes sp.* | 37.00 | 800.00 | 22.26 |
| 2019 | Breeding | Berlengas | *Atherina boyeri* | 4.14 | 12.86 | 1.15 |
| 2019 | Breeding | Berlengas | Atherina presbyter | 2.29 | 11.43 | 0.72 |
| 2019 | Breeding | Berlengas | *Micromesistius poutassou* | 0.14 | 1.43 | |
| 2019 | Breeding | Berlengas | *Trisopterus luscus* | 1.29 | 12.86 | 3.97 |

| Year | Period | Area | Species | FN | FO | FB |
|---|---|---|---|---|---|---|
| 2019 | Breeding | Berlengas | *Trisopterus minutus* | 2.43 | 18.57 | 5.96 |
| 2019 | Breeding | Berlengas | *Gobius paganellus* | 0.29 | 2.86 | 0.09 |
| 2019 | Breeding | Berlengas | *Coris julis* | 13.57 | 57.14 | 14.58 |
| 2019 | Breeding | Berlengas | *Ctenolabrus rupestris* | 0.14 | 1.43 | 0.03 |
| 2019 | Breeding | Berlengas | *Labrus bergylta* | 0.86 | 7.14 | 2.49 |
| 2019 | Breeding | Berlengas | *Labrus mixtus* | 7.14 | 42.86 | 7.09 |
| 2019 | Breeding | Berlengas | *Labrus viridis* | 2.00 | 18.57 | 0.74 |
| 2019 | Breeding | Berlengas | *Symphodus bailloni* | 2.29 | 21.43 | 1.93 |
| 2019 | Breeding | Berlengas | *Symphodus cinereus* | 3.29 | 24.29 | 1.71 |
| 2019 | Breeding | Berlengas | *Symphodus melops* | 2.29 | 20.00 | 1.58 |
| 2019 | Breeding | Berlengas | *Symphodus trutta* | 0.43 | 4.29 | 0.24 |
| 2019 | Breeding | Berlengas | *Thalassoma pavo* | 0.14 | 1.43 | 1.51 |
| 2019 | Breeding | Berlengas | Coryphaenoides mediterraneus | 0.14 | 1.43 | 0.27 |
| 2019 | Breeding | Berlengas | *Antimora rostrata* | 0.14 | 1.43 | 1.17 |
| 2019 | Breeding | Berlengas | *Gaidropsarus mediterraneus* | 1.86 | 15.71 | 4.53 |
| 2019 | Breeding | Berlengas | *Serranus cabrilla* | 1.57 | 14.29 | 2.60 |
| 2019 | Breeding | Berlengas | *Pegusa lascaris* | 0.86 | 4.29 | 3.00 |
| 2019 | Breeding | Berlengas | *Boops boops* | 0.43 | 2.86 | 0.29 |
| 2019 | Breeding | Berlengas | *Dentex dentex* | 0.14 | 1.43 | 0.27 |
| 2019 | Breeding | Berlengas | *Labrus merula* | 1.57 | 15.71 | 11.97 |
| 2019 | Breeding | Berlengas | *Diplodus sargus* | 0.43 | 4.29 | 1.02 |
| 2019 | Breeding | Berlengas | *Diplodus vulgaris* | 0.14 | 1.43 | 0.97 |
| 2019 | Breeding | Berlengas | *Pagellus bogaraveo* | 0.14 | 1.43 | 0.03 |
| 2019 | Breeding | Berlengas | *Sparus aurata* | 0.29 | 2.86 | 0.00 |
| 2019 | Breeding | Berlengas | *Spondyliosoma cantharus* | 0.29 | 1.43 | 0.80 |
| 2019 | Breeding | Berlengas | *Trigla lyra* | 0.43 | 4.29 | 7.26 |
| 2019 | Non-Breeding | Berlengas | *Ammodytes sp.* | 11.76 | 33.33 | 17.40 |
| 2019 | Non-Breeding | Berlengas | *Trisopterus luscus* | 5.88 | 33.33 | |
| 2019 | Non-Breeding | Berlengas | *Labrus mixtus* | 52.94 | 66.67 | |
| 2019 | Non-Breeding | Berlengas | *Symphodus cinereus* | 5.88 | 33.33 | 21.59 |
| 2019 | Non-Breeding | Berlengas | *Serranus cabrilla* | 5.88 | 33.33 | 61.01 |
| 2020 | Breeding | Berlengas | *Ammodytes sp.* | 25.93 | 53.33 | 19.64 |
| 2020 | Breeding | Berlengas | *Ammodytidae* | 2.47 | 20.00 | 1.86 |
| 2020 | Breeding | Berlengas | *Atherina presbyter* | 27.78 | 13.33 | 5.23 |
| 2020 | Breeding | Berlengas | *Bothidae* | 1.85 | 6.67 | |
| 2020 | Breeding | Berlengas | *Trisopterus luscus* | 4.32 | 26.67 | 2.88 |
| 2020 | Breeding | Berlengas | *Trisopterus minutus* | 7.41 | 26.67 | 17.44 |
| 2020 | Breeding | Berlengas | *Gobius paganellus* | 0.62 | 6.67 | 0.26 |
| 2020 | Breeding | Berlengas | *Coris julis* | 1.85 | 20.00 | 16.79 |
| 2020 | Breeding | Berlengas | *Ctenolabrus rupestris* | 0.62 | 6.67 | 1.95 |
| 2020 | Breeding | Berlengas | *Labrus bergylta* | 3.09 | 20.00 | 10.19 |
| 2020 | Breeding | Berlengas | *Labrus mixtus* | 1.85 | 20.00 | |
| 2020 | Breeding | Berlengas | *Labrus viridis* | 0.62 | 6.67 | 0.11 |
| 2020 | Breeding | Berlengas | *Symphodus mediterraneus* | 1.85 | 6.67 | 1.34 |
| 2020 | Breeding | Berlengas | *Symphodus melops* | 0.62 | 6.67 | 0.35 |
| 2020 | Breeding | Berlengas | *Symphodus roissali* | 1.23 | 6.67 | 1.13 |
| 2020 | Breeding | Berlengas | *Symphodus sp.* | 0.62 | 6.67 | 0.35 |
| 2020 | Breeding | Berlengas | *Gaidropsarus mediterraneus* | 0.62 | 6.67 | 0.53 |
| 2020 | Breeding | Berlengas | *Serranus cabrilla* | 1.85 | 20.00 | 4.10 |

| Year | Period | Area | Species | FN | FO | FB |
|---|---|---|---|---|---|---|
| 2020 | Breeding | Berlengas | *Pegusa lascaris* | 1.23 | 6.67 | 0.30 |
| 2020 | Breeding | Berlengas | *Boops boops* | 0.62 | 6.67 | 0.52 |
| 2020 | Breeding | Berlengas | *Dentex dentex* | 0.62 | 6.67 | 0.02 |
| 2020 | Breeding | Berlengas | *Diplodus vulgaris* | 1.23 | 6.67 | 0.19 |
| 2020 | Breeding | Berlengas | *Diplodus annularis* | 1.23 | 6.67 | 1.17 |
| 2020 | Breeding | Berlengas | *Diplodus sp.* | 1.23 | 6.67 | 1.34 |
| 2020 | Non-Breeding | Berlengas | *Ammodytidae* | 36.96 | 42.86 | 0.12 |
| 2020 | Non-Breeding | Berlengas | *Atherina presbyter* | 2.17 | 14.29 | 1.11 |
| 2020 | Non-Breeding | Berlengas | *Parablennius sp.* | 2.17 | 14.29 | 0.00 |
| 2020 | Non-Breeding | Berlengas | *Trisopterus minutus* | 17.39 | 14.29 | 0.37 |
| 2020 | Non-Breeding | Berlengas | *Gobius sp.* | 2.17 | 14.29 | 0.00 |
| 2020 | Non-Breeding | Berlengas | *Coris julis* | 2.17 | 14.29 | 0.07 |
| 2020 | Non-Breeding | Berlengas | *Ctenolabrus rupestris* | 6.52 | 14.29 | 0.02 |
| 2020 | Non-Breeding | Berlengas | *Labrus merula* | 2.17 | 14.29 | 0.09 |
| 2020 | Non-Breeding | Berlengas | *Labrus mixtus* | 2.17 | 14.29 | 0.02 |
| 2020 | Non-Breeding | Berlengas | *Symphodus cinereus* | 2.17 | 14.29 | 0.01 |
| 2020 | Non-Breeding | Berlengas | *Symphodus melops* | 4.35 | 14.29 | 0.10 |
| 2020 | Non-Breeding | Berlengas | *Spondyliosoma cantharus* | 4.35 | 14.29 | 0.01 |
| 2020 | Non-Breeding | Arrabida | *Ammodytes sp.* | 0.91 | 4.55 | 0.40 |
| 2020 | Non-Breeding | Arrabida | *Ammodytes tobianus* | 5.45 | 11.36 | 6.44 |
| 2020 | Non-Breeding | Arrabida | *Hyperoplus lanceolatus* | 26.67 | 34.09 | 42.02 |
| 2020 | Non-Breeding | Arrabida | *Ammodytidae* | 23.03 | 25.00 | 19.94 |
| 2020 | Non-Breeding | Arrabida | *Atherina boyeri* | 0.30 | 2.27 | 0.13 |
| 2020 | Non-Breeding | Arrabida | *Atherina presbyter* | 5.15 | 2.27 | 1.21 |
| 2020 | Non-Breeding | Arrabida | *Atherina sp.* | 0.91 | 2.27 | 1.21 |
| 2020 | Non-Breeding | Arrabida | *Parablennius rouxi* | 0.61 | 2.27 | |
| 2020 | Non-Breeding | Arrabida | *Callionymus maculatus* | 0.61 | 2.27 | |
| 2020 | Non-Breeding | Arrabida | *Trisopterus minutus* | 1.21 | 4.55 | 1.27 |
| 2020 | Non-Breeding | Arrabida | *Gobius paganellus* | 0.61 | 4.55 | 0.14 |
| 2020 | Non-Breeding | Arrabida | *Coris julis* | 6.36 | 18.18 | 9.02 |
| 2020 | Non-Breeding | Arrabida | *Ctenolabrus rupestris* | 0.61 | 2.27 | 1.10 |
| 2020 | Non-Breeding | Arrabida | *Labridae* | 0.30 | 2.27 | 42.02 |
| 2020 | Non-Breeding | Arrabida | *Labrus bergylta* | 0.61 | 2.27 | 3.14 |
| 2020 | Non-Breeding | Arrabida | *Labrus mixtus* | 2.73 | 9.09 | 3.65 |
| 2020 | Non-Breeding | Arrabida | *Symphodus bailloni* | 0.61 | 4.55 | 0.44 |
| 2020 | Non-Breeding | Arrabida | *Symphodus cinereus* | 0.61 | 4.55 | 1.46 |
| 2020 | Non-Breeding | Arrabida | *Symphodus melops* | 0.61 | 4.55 | 1.95 |
| 2020 | Non-Breeding | Arrabida | *Phycidae* | 0.30 | 2.27 | |
| 2020 | Non-Breeding | Arrabida | *Serranus cabrilla* | 0.61 | 2.27 | 2.68 |
| 2020 | Non-Breeding | Arrabida | *Serranus hepatus* | 0.61 | 2.27 | 0.04 |
| 2020 | Non-Breeding | Arrabida | *Soleidae* | 0.91 | 6.82 | |
| 2020 | Non-Breeding | Arrabida | *Boops boops* | 0.91 | 4.55 | 0.94 |
| 2020 | Non-Breeding | Arrabida | *Sarpa salpa* | 0.30 | 2.27 | |
| 2020 | Non-Breeding | Arrabida | *Sparidae* | 3.64 | 2.27 | |
| 2020 | Non-Breeding | Arrabida | *Trachinus draco* | 0.30 | 2.27 | 0.02 |
| 2020 | Non-Breeding | Arrabida | *Arnoglossus imperialis* | 0.91 | 6.82 | 2.60 |
| 2021 | Breeding | Berlengas | *Ammodytidae* | 36.09 | 28.00 | 3.77 |
| 2021 | Breeding | Berlengas | *Atherina presbyter* | 2.37 | 4.00 | |
| 2021 | Breeding | Berlengas | *Atherina sp.* | 0.59 | 4.00 | |

| Year | Period | Area | Species | FN | FO | FB |
|---|---|---|---|---|---|---|
| 2021 | Breeding | Berlengas | *Parablennius sp.* | 1.18 | 8.00 | 0.24 |
| 2021 | Breeding | Berlengas | *Sprattus sprattus* | 0.59 | 4.00 | |
| 2021 | Breeding | Berlengas | *Clupeidae* | 1.18 | 4.00 | |
| 2021 | Breeding | Berlengas | *Trisopterus luscus* | 2.96 | 20.00 | 1.64 |
| 2021 | Breeding | Berlengas | *Trisopterus sp.* | 1.18 | 8.00 | |
| 2021 | Breeding | Berlengas | *Gadidae* | 0.59 | 4.00 | |
| 2021 | Breeding | Berlengas | *Gobius sp.* | 0.59 | 4.00 | 0.18 |
| 2021 | Breeding | Berlengas | *Acantholabrus palloni* | 0.59 | 4.00 | |
| 2021 | Breeding | Berlengas | *Coris julis* | 3.55 | 20.00 | 4.24 |
| 2021 | Breeding | Berlengas | *Ctenolabrus rupestris* | 5.33 | 28.00 | 0.99 |
| 2021 | Breeding | Berlengas | *Symphodus sp.* | 5.92 | 28.00 | 1.31 |
| 2021 | Breeding | Berlengas | *Symphodus bailloni* | 1.18 | 8.00 | |
| 2021 | Breeding | Berlengas | *Symphodus cinereus* | 0.59 | 4.00 | |
| 2021 | Breeding | Berlengas | *Symphodus mediterraneus* | 0.59 | 4.00 | |
| 2021 | Breeding | Berlengas | *Symphodus melops* | 0.59 | 4.00 | |
| 2021 | Breeding | Berlengas | *Labrus merula* | 0.59 | 4.00 | 2.44 |
| 2021 | Breeding | Berlengas | *Labrus mixtus* | 2.37 | 12.00 | 0.39 |
| 2021 | Breeding | Berlengas | *Labrus sp.* | 1.18 | 8.00 | |
| 2021 | Breeding | Berlengas | *Thalassoma pavo* | 0.59 | 4.00 | 83.61 |
| 2021 | Breeding | Berlengas | *Labridae* | 5.92 | 24.00 | |
| 2021 | Breeding | Berlengas | *Moronidae* | 0.59 | 4.00 | |
| 2021 | Breeding | Berlengas | *Ciliata mustela* | 2.96 | 16.00 | |
| 2021 | Breeding | Berlengas | *Physicidae* | 0.59 | 4.00 | 0.45 |
| 2021 | Breeding | Berlengas | *Mycteroperca sp.* | 1.18 | 4.00 | |
| 2021 | Breeding | Berlengas | *Serranus sp.* | 4.14 | 12.00 | |
| 2021 | Breeding | Berlengas | *Serranidae* | 0.59 | 4.00 | |
| 2021 | Breeding | Berlengas | *Dentex sp.* | 0.59 | 4.00 | |
| 2021 | Breeding | Berlengas | *Diplodus vulgaris* | 0.59 | 4.00 | 0.32 |
| 2021 | Breeding | Berlengas | *Sparidae* | 4.14 | 24.00 | |
| 2021 | Breeding | Berlengas | *Trachinus draco* | 1.18 | 4.00 | 0.43 |
| 2021 | Non-Breeding | Berlengas | *Ammodytidae* | 85.71 | 100 | 7.27 |
| 2021 | Non-Breeding | Berlengas | *Gadidae* | 4.76 | 50 | |
| 2021 | Non-Breeding | Berlengas | *Labrus merula* | 4.76 | 50 | |
| 2021 | Non-Breeding | Berlengas | *Sparidae* | 4.76 | 50 | |
| 2021 | Breeding | Arrabida | *Ammodytes tobianus* | 9.41 | 22 | 3.09 |
| 2021 | Breeding | Arrabida | *Hyperoplus lanceolatus* | 21.44 | 42 | 5.56 |
| 2021 | Breeding | Arrabida | *Ammodytidae* | 26.69 | 32 | |
| 2021 | Breeding | Arrabida | *Atherina boyeri* | 1.94 | 6 | 0.19 |
| 2021 | Breeding | Arrabida | *Atherina presbyter* | 5.12 | 6 | 0.37 |
| 2021 | Breeding | Arrabida | *Atherina sp.* | 0.14 | 2 | |
| 2021 | Breeding | Arrabida | *Parablennius gattorugine* | 0.55 | 6 | |
| 2021 | Breeding | Arrabida | *Parablennius pilicornis* | 0.14 | 2 | 0.04 |
| 2021 | Breeding | Arrabida | *Parablennius rouxi* | 0.14 | 2 | |
| 2021 | Breeding | Arrabida | *Arnoglossus imperialis* | 0.55 | 6 | 0.25 |
| 2021 | Breeding | Arrabida | Arnoglossus rueppelii | 0.14 | 2 | |
| 2021 | Breeding | Arrabida | *Callionymus maculatus* | 0.41 | 6 | |
| 2021 | Breeding | Arrabida | *Trisopterus luscus* | 1.66 | 2 | 0.08 |
| 2021 | Breeding | Arrabida | *Trisopterus minutus* | 4.56 | 4 | 0.12 |
| 2021 | Breeding | Arrabida | *Aphia minuta* | 2.21 | 14 | |

| Year | Period | Area | Species | FN | FO | FB |
|---|---|---|---|---|---|---|
| 2021 | Breeding | Arrabida | *Coris julis* | 1.94 | 18 | 1.03 |
| 2021 | Breeding | Arrabida | *Ctenolabrus rupestris* | 1.24 | 12 | 0.49 |
| 2021 | Breeding | Arrabida | *Labrus bergylta* | 0.14 | 2 | 0.33 |
| 2021 | Breeding | Arrabida | *Labrus mixtus* | 0.83 | 4 | 0.26 |
| 2021 | Breeding | Arrabida | *Symphodus bailloni* | 0.14 | 2 | 0.01 |
| 2021 | Breeding | Arrabida | *Symphodus cinereus* | 0.69 | 8 | 0.11 |
| 2021 | Breeding | Arrabida | *Symphodus melops* | 0.28 | 4 | 0.04 |
| 2021 | Breeding | Arrabida | *Moronidae* | 0.28 | 2 | |
| 2021 | Breeding | Arrabida | *Ciliata mustela* | 0.41 | 6 | |
| 2021 | Breeding | Arrabida | *Pegusa lascaris* | 0.14 | 2 | 0.06 |
| 2021 | Breeding | Arrabida | *Soleidae* | 4.84 | 12 | 0.10 |
| 2021 | Breeding | Arrabida | *Boops boops* | 0.28 | 2 | 0.35 |
| 2021 | Breeding | Arrabida | *Gobius paganellus* | 0.41 | 6 | 0.10 |
| 2021 | Breeding | Arrabida | *Diplodus bellotti* | 1.11 | 2 | |
| 2021 | Breeding | Arrabida | *Gobius sp.* | 2.77 | 2 | 0.03 |
| 2021 | Breeding | Arrabida | *Diplodus sp.* | 0.28 | 2 | 0.03 |
| 2021 | Breeding | Arrabida | *Pagellus bogaraveo* | 0.14 | 2 | 0.01 |
| 2021 | Breeding | Arrabida | *Spondyliosoma cantharus* | 0.69 | 2 | 0.22 |
| 2022 | Breeding | Berlengas | *Ammodytidae* | 28.67 | 23.33 | |
| 2022 | Breeding | Berlengas | *Trisopterus luscus* | 6.00 | 20.00 | |
| 2022 | Breeding | Berlengas | *Trisopterus sp.* | 14.67 | 30.00 | |
| 2022 | Breeding | Berlengas | *Gobiidae* | 0.67 | 3.33 | |
| 2022 | Breeding | Berlengas | *Ctenolabrus rupestris* | 0.67 | 3.33 | |
| 2022 | Breeding | Berlengas | *Labrus mixtus* | 0.67 | 3.33 | |
| 2022 | Breeding | Berlengas | *Labrus sp.* | 0.67 | 3.33 | |
| 2022 | Breeding | Berlengas | *Symphodus sp.* | 3.33 | 10.00 | |
| 2022 | Breeding | Berlengas | *Labridae* | 24.67 | 63.33 | |
| 2022 | Breeding | Berlengas | *Serranus sp.* | 4.00 | 10.00 | |
| 2022 | Breeding | Berlengas | *Serranidae* | 2.67 | 10.00 | |
| 2022 | Breeding | Berlengas | *Sparidae* | 0.67 | 3.33 | |
| 2022 | Non-Breeding | Berlengas | *Ammodytidae* | 17.24 | 33.33 | |
| 2022 | Non-Breeding | Berlengas | *Trisopterus luscus* | 13.79 | 33.33 | |
| 2022 | Non-Breeding | Berlengas | *Trisopterus sp.* | 6.90 | 16.67 | |
| 2022 | Non-Breeding | Berlengas | *Gadidae* | 17.24 | 33.33 | |
| 2022 | Non-Breeding | Berlengas | *Labridae* | 6.90 | 33.33 | |
| 2022 | Non-Breeding | Berlengas | *Serranus sp.* | 3.45 | 16.67 | |
| 2022 | Non-Breeding | Berlengas | *Sparidae* | 6.90 | 16.67 | |
| 2023 | Breeding | Berlengas | *Ammodytidae* | 55.93 | 35.59 | |
| 2023 | Breeding | Berlengas | *Trisopterus sp.* | 3.60 | 16.95 | |
| 2023 | Breeding | Berlengas | *Gadidae* | 0.35 | 5.08 | |
| 2023 | Breeding | Berlengas | *Gobius sp.* | 0.47 | 5.08 | |
| 2023 | Breeding | Berlengas | *Gobiidae* | 0.58 | 6.78 | |
| 2023 | Breeding | Berlengas | *Coris julis* | 2.21 | 23.73 | |
| 2023 | Breeding | Berlengas | *Ctenolabrus rupestris* | 0.23 | 3.39 | |
| 2023 | Breeding | Berlengas | *Labrus bergylta* | 0.12 | 1.69 | |
| 2023 | Breeding | Berlengas | *Labrus sp.* | 1.40 | 13.56 | |
| 2023 | Breeding | Berlengas | *Symphodus bailloni* | 0.23 | 3.39 | |
| 2023 | Breeding | Berlengas | *Symphodus cinereus* | 0.35 | 5.08 | |
| 2023 | Breeding | Berlengas | *Symphodus mediterraneus* | 0.70 | 3.39 | |

| Year | Period | Area | Species | FN | FO | FB |
|---|---|---|---|---|---|---|
| 2023 | Breeding | Berlengas | *Symphodus melops* | 0.58 | 8.47 | |
| 2023 | Breeding | Berlengas | *Symphodus roissali* | 0.12 | 1.69 | |
| 2023 | Breeding | Berlengas | *Symphodus sp.* | 3.49 | 30.51 | |
| 2023 | Breeding | Berlengas | *Labridae* | 9.65 | 49.15 | |
| 2023 | Breeding | Berlengas | *Lotidae* | 0.35 | 3.39 | |
| 2023 | Breeding | Berlengas | *Dicentrarchus sp.* | 0.35 | 3.39 | |
| 2023 | Breeding | Berlengas | *Ciliata mustela* | 0.58 | 6.78 | |
| 2023 | Breeding | Berlengas | *Phycis sp.* | 0.23 | 1.69 | |
| 2023 | Breeding | Berlengas | *Phycidae* | 0.58 | 8.47 | |
| 2023 | Breeding | Berlengas | *Mycteroperca fusca* | 0.12 | 1.69 | |
| 2023 | Breeding | Berlengas | *Serranus cabrilla* | 1.16 | 10.17 | |
| 2023 | Breeding | Berlengas | *Serranus sp.* | 0.23 | 3.39 | |
| 2023 | Breeding | Berlengas | *Serranidae* | 0.70 | 10.17 | |
| 2023 | Breeding | Berlengas | *Soleidae* | 0.58 | 1.69 | |
| 2023 | Breeding | Berlengas | *Diplodus annularis* | 0.12 | 1.69 | |
| 2023 | Breeding | Berlengas | *Trisopterus minutus* | 0.12 | 1.69 | |
| 2023 | Breeding | Berlengas | *Trisopterus luscus* | 5.70 | 40.68 | |
| 2023 | Breeding | Berlengas | *Gobius paganellus* | 0.35 | 3.39 | |
| 2023 | Breeding | Berlengas | *Diplodus vulgaris* | 0.12 | 1.69 | |
| 2023 | Breeding | Berlengas | *Diplodus sp.* | 0.23 | 1.69 | |
| 2023 | Breeding | Berlengas | *Sparidae* | 1.28 | 11.86 | |
| 2023 | Breeding | Berlengas | *Tranchinidae* | 0.12 | 1.69 | |
| 2023 | Breeding | Berlengas | *Trigla lyra* | 0.12 | 1.69 | |
| 2023 | Non-Breeding | Berlengas | Labridae | 71.43 | 100 | |
| 2024 | Breeding | Berlengas | *Ammodytes tobianus* | 7.36 | 37.04 | 0.54 |
| 2024 | Breeding | Berlengas | *Hyperoplus lanceolatus* | 0.92 | 11.11 | 0.15 |
| 2024 | Breeding | Berlengas | *Atherina boyeri* | 3.68 | 7.41 | 0.09 |
| 2024 | Breeding | Berlengas | *Parablennius gattorugine* | 13.19 | 22.22 | |
| 2024 | Breeding | Berlengas | *Parablennius pilicornis* | 0.61 | 3.70 | |
| 2024 | Breeding | Berlengas | *Parablennius rouxi* | 3.07 | 25.93 | |
| 2024 | Breeding | Berlengas | *Arnoglossus imperialis* | 0.92 | 7.41 | 0.03 |
| 2024 | Breeding | Berlengas | *Arnoglossus rueppelii* | 4.29 | 29.63 | |
| 2024 | Breeding | Berlengas | *Arnoglossus thori* | 0.92 | 14.81 | |
| 2024 | Breeding | Berlengas | *Bothus podas* | 0.31 | 3.70 | |
| 2024 | Breeding | Berlengas | *Callionymus lyra* | 1.84 | 11.11 | |
| 2024 | Breeding | Berlengas | *Callionymus maculatus* | 0.31 | 3.70 | |
| 2024 | Breeding | Berlengas | *Trachurus trachurus* | 0.31 | 3.70 | 0.02 |
| 2024 | Breeding | Berlengas | *Engraulis encrasicolus* | 0.61 | 7.41 | 0.63 |
| 2024 | Breeding | Berlengas | *Trisopterus luscus* | 2.45 | 22.22 | 1.64 |
| 2024 | Breeding | Berlengas | *Trisopterus minutus* | 1.84 | 11.11 | 0.63 |
| 2024 | Breeding | Berlengas | *Aphia minuta* | 1.84 | 22.22 | |
| 2024 | Breeding | Berlengas | *Coris julis* | 9.51 | 33.33 | 0.26 |
| 2024 | Breeding | Berlengas | *Gobius paganellus* | 1.23 | 11.11 | 0.00 |
| 2024 | Breeding | Berlengas | *Ctenolabrus rupestris* | 0.61 | 7.41 | 0.05 |
| 2024 | Breeding | Berlengas | *Labrus mixtus* | 1.53 | 14.81 | 0.17 |
| 2024 | Breeding | Berlengas | *Symphodus bailloni* | 1.84 | 22.22 | 0.19 |
| 2024 | Breeding | Berlengas | *Symphodus cinereus* | 3.99 | 29.63 | 0.26 |
| 2024 | Breeding | Berlengas | *Symphodus melops* | 1.84 | 22.22 | 0.12 |
| 2024 | Breeding | Berlengas | *Thalassoma pavo* | 0.61 | 7.41 | |

| Year | Period | Area | Species | FN | FO | FB |
|---|---|---|---|---|---|---|
| 2024 | Breeding | Berlengas | *Dicentrarchus sp.* | 0.92 | 7.41 | |
| 2024 | Breeding | Berlengas | *Mugil cephalus* | 0.31 | 3.70 | |
| 2024 | Breeding | Berlengas | *Ciliata mustela* | 1.53 | 18.52 | 94.68 |
| 2024 | Breeding | Berlengas | *Helicolenus dactylopterus* | 0.31 | 3.70 | |
| 2024 | Breeding | Berlengas | *Serranus cabrilla* | 0.31 | 3.70 | 0.05 |
| 2024 | Breeding | Berlengas | *Serranus hepatus* | 0.31 | 3.70 | |
| 2024 | Breeding | Berlengas | *Microchirus boscanion* | 2.45 | 7.41 | |
| 2024 | Breeding | Berlengas | *Boops boops* | 0.61 | 7.41 | 0.41 |
| 2024 | Breeding | Berlengas | *Diplodus annularis* | 0.31 | 3.70 | |
| 2024 | Breeding | Berlengas | *Diplodus bellotti* | 1.84 | 11.11 | |
| 2024 | Breeding | Berlengas | *Diplodus cervinus* | 0.31 | 3.70 | |
| 2024 | Breeding | Berlengas | *Diplodus sargus* | 1.53 | 11.11 | 0.01 |
| 2024 | Breeding | Berlengas | *Diplodus vulgaris* | 0.31 | 3.70 | |
| 2024 | Breeding | Berlengas | *Diplodus sp.* | 3.37 | 7.41 | 0.08 |
| 2024 | Breeding | Berlengas | *Sarpa salpa* | 15.64 | 7.41 | |
| 2024 | Breeding | Berlengas | *Spondyliosoma cantharus* | 0.31 | 3.70 | |
| 2024 | Breeding | Berlengas | *Syngnathus acus* | 0.31 | 3.70 | |

*Table S3. Top 10 prey species selected by Maximum IRI (2016-2024). "MAX IRI (Year)" indicates the highest IRI observed for the species and the year in which it occurred. "Mean IRI± SD" shows the average IRI and standard deviation across 2016-2024. FN%Max, FO% Max and FB%Max correspond, respectively, to the numerical frequency, frequency of occurrence, and biomass percentage in the year of Maximum IRI. When marked as "NA," data were not available for that year. "Years (%Present)" indicates the number of years in which the species was recorded and the percentage of the total period*

| Prey species | Max IRI (Year) | Mean IRI ± SD | FN% Max | FO% Max | FB% Max | Years (%Present) |
|---|---|---|---|---|---|---|
| **Ammodytidae** | 47411.72 (2019) | 4169.05 ± 10802.72 | 37.00 | 80.00 | 22.26 | 9 (100%) |
| **Atherinidae** *Atherina presbyter* | 440.16 (2020) | 82.67 ± 158.47 | 27.78 | 13.33 | 5.23 | 4 (44.4%) |
| **Carangidae** *Trachurus trachurus* | 1662.72 (2018) | 563.05 ± 952.42 | 6.98 | 22.22 | 67.85 | 3 (33.3%) |
| **Gadidae** *Trisopterus luscus* | 10616.20 (2016) | 1562.34 ± 3559.65 | 47.27 | 75.00 | 94.28 | 9 (100%) |
| **Labridae** *Coris julis* | 3653.88 (2016) | 808.24 ± 1016.19 | 33.33 | 40.00 | 58.01 | 8 (88.9%) |
| *Labrus mixtus* | 3529.41 (2019) | 340.76 ± 971.67 | 52.94 | 66.67 | NA | 8 (88.9%) |
| *Symphodus bailloni* | 1385.18 (2016) | 215.45 ± 433.02 | 5.45 | 37.50 | 31.48 | 8 (88.9%) |
| **Phycidae** *Ciliata Mustela* | 1812.86 (2016) | 248.09 ± 632.66 | 8.33 | 40.00 | 36.99 | 5 (55.6%) |
| *Gaidropsarus mediterraneus* | 6794.45 (2018) | 1726.33 ± 3379.05 | 36.36 | 100.00 | 31.58 | 4 (44.44%) |
| **Serranidae** *Serranus cabrilla* | 2634.43 (2016) | 606.13 ± 1009.22 | 8.33 | 40.00 | 57.53 | 7 (77.8%) |
